# Unipolar peptidoglycan synthesis in the Rhizobiales requires an essential class A penicillin-binding protein

**DOI:** 10.1101/2021.03.31.437934

**Authors:** Michelle A. Williams, Alena Aliashkevich, Elizaveta Krol, Erkin Kuru, Jacob M. Bouchier, Jonathan Rittichier, Yves V. Brun, Michael S. VanNieuwenhze, Anke Becker, Felipe Cava, Pamela J. B. Brown

## Abstract

Members of the Rhizobiales are polarly-growing bacteria that lack homologs of the canonical rod complex. To investigate the mechanisms underlying polar cell wall synthesis, we systematically probed the function of cell wall synthesis enzymes in the plant-pathogen *Agrobacterium tumefaciens*. The development of fluorescent d-amino acid dipeptide (FDAAD) probes, which are incorporated into peptidoglycan by penicillin-binding proteins in *A. tumefaciens*, enabled us to monitor changes in growth patterns in the mutants. Use of these fluorescent cell wall probes and peptidoglycan compositional analysis convincingly demonstrate that a single class A penicillin-binding protein is essential for polar peptidoglycan synthesis. Furthermore, we find evidence of an alternative mode of cell wall synthesis that likely requires ld-transpeptidase activity. Genetic analysis and cell wall targeting antibiotics reveal that the mechanism of unipolar growth is conserved in *Sinorhizobium* and *Brucella*. This work provides insights into unipolar peptidoglycan biosynthesis employed by the Rhizobiales during cell elongation.

## INTRODUCTION

Our current understanding of peptidoglycan (PG) assembly in rod-shaped bacteria stems largely from investigations conducted using well-known model species like *Escherichia coli* and *Bacillus subtilis*, which incorporate new cell wall material along the lateral sidewalls of the cell body. Expanding our studies of cell wall synthesis to include diverse species, with alternative modes of elongation, is an important step in unveiling the mechanisms of how and why bacteria evolve novel growth modes and generate innovative morphologies. It had, for example, long been assumed that all rod-shaped bacteria employed the same growth strategy; however, unipolar growth is widespread among rod-shaped bacteria in the Alphaproteobacterial order Rhizobiales, suggesting diversification of growth strategies [1]. The Rhizobiales are comprised of diverse bacteria with respect to both their cellular morphology, and their environmental niches [2, 3]. This includes many species of medical and agricultural significance such as the facultative intracellular pathogens *Bartonella* and *Brucella*, the nitrogen-fixing plant symbiont *Sinorhizobium*, and the causative agent of crown gall disease *Agrobacterium tumefaciens*, [4, 5]. PG labeling experiments in *A. tumefaciens, Brucella abortus*, and *S. meliloti* have confirmed that unipolar growth is the mode of elongation utilized by these rod-shaped species [1]. Remarkably, labeling experiments have revealed that polar growth in *A. tumefaciens* results specifically in the incorporation of pentapeptides at the growth tip; [6] however, the mechanisms that underlie polar PG biosynthesis remain poorly understood.

PG biosynthesis is an essential process that allows bacteria to grow and divide, faithfully reproducing their characteristic cell shape [7]. PG assembly requires different classes of synthesis enzymes including the penicillin-binding proteins (PBPs), which can be further divided into two classes. Class A PBPs are bifunctional enzymes that catalyze β-1,4 linkages between the N-Acetylglucosamine (NAG) and N-Acetylmuramic acid (NAM) sugars in a process called transglycosylation, and also synthesize crosslinks between 4-3 and 4-4 peptides in a process known as transpeptidation [8]. The class B PBPs are monofunctional enzymes that have only transpeptidase (TPase) activity [9]. The shape, elongation, division, sporulation (SEDS) family proteins, RodA and FtsW, also possess glycosyltransferase (GTase) activity [10, 11]. Current models of cell wall assembly maintain that SEDS proteins, in complex with their cognate class B PBP, are the primary drivers of cell wall synthesis and are required to sustain rod shape [12–14]. Thus, RodA functions with PBP2 during elongation, while FtsW functions with PBP3 (encoded by *ftsI*) during cell division. The class A PBPs are currently thought to act independently from the rod complex, functioning primarily in PG maintenance and repair [15, 16].

The suite of cell wall synthesis enzymes encoded by the Rhizobiales, is distinctly different from other bacterial orders. For example, the elongation-specific rod complex of proteins including PBP2, RodA, and MreBCD are universally absent [1, 17], suggesting that RodA-PBP2 are not the primary drivers of elongation. In addition, the genomes of Rhizobiales are enriched for the presence of ld-transpeptidases (LDTs). LDTs are a class of cell wall synthesizing enzymes that carry out transpeptidation reactions to catalyze 3-4 and 3-3 crosslinks in the cell wall [18]. The cell wall of *A. tumefaciens* contains a high proportion (30%) of 3-3 and 3-4 crosslinks, in contrast to laterally growing rod-shaped bacteria, where only 1-5% of the cell wall is crosslinked by LDTs [19]. This suggests that LDT enzymes may play an important role during polar growth [1, 20]. Overall, these observations suggest that Rhizobiales use a non-canonical mechanism for polar elongation.

Using a combination of microscopy, new probes, biochemical, and genetic analyses we have characterized the function of the six high molecular weight PBPs encoded by *A. tumefaciens* and have identified the major cell wall synthesis enzymes required for polar growth. We show that, unlike the proposed auxiliary function of PBP1a in other rod-shaped bacteria, in *A. tumefaciens* PBP1a is an essential enzyme required for polar PG expansion, with depletion of PBP1a resulting in a loss of proper rod-shape. Using newly developed fluorescent d-amino acid dipeptide (FDAAD) probes, we show that PBP1a is the enzyme primarily responsible for inserting nascent PG at the pole. Additionally, PBP1a depletion leads to a modified PG composition, including an increase in LDT linkages. Collectively, this suggests that the mechanism of polar growth in the Rhizobiales has evolved through the expansion, diversification, and altered regulation of the core cell wall synthesis machinery. We confirmed the essentiality of PBP1a in the closely related bacterium *Sinorhizobium*, suggesting that the mechanisms underlying polar growth in the Rhizobiales are well conserved. Finally, we have identified the β-lactam faropenem as a specific inhibitor of polar growth in *Agrobacterium, Sinorhizobium*, and *Brucella*, indicating that the target(s) of faropenem are conserved among the Rhizobiales. These findings broaden our understanding of the role of PG synthesis enzymes that contribute to polar growth and will inform strategies aimed at developing novel therapeutics that target the cell wall of polar-growing bacteria in the Rhizobiales [21, 22].

## RESULTS

### The class A cell wall synthase PBP1a is essential for polar growth

To begin probing the molecular determinant(s) of polar growth in *A. tumefaciens*, we sought to generate deletions of the predicted PG synthase enzymes. Although *A. tumefaciens* lacks homologs to the predicted cell elongation synthases RodA and PBP2, the genome encodes four bifunctional class A PBPs (PBP1a, PBP1b1, PBP1b2 and PBP1c), two monofunctional class B PBPs (PBP3a and PBP3b), and one monofunctional GTase, MtgA (Figures 1A, 2A). To determine which of these enzyme(s) provide the primary GTase activity in the absence of a RodA homolog, we made in-frame deletions of those genes encoding predicted GTase enzymes to further explore their contribution during cell growth or division.

**Figure 1.**
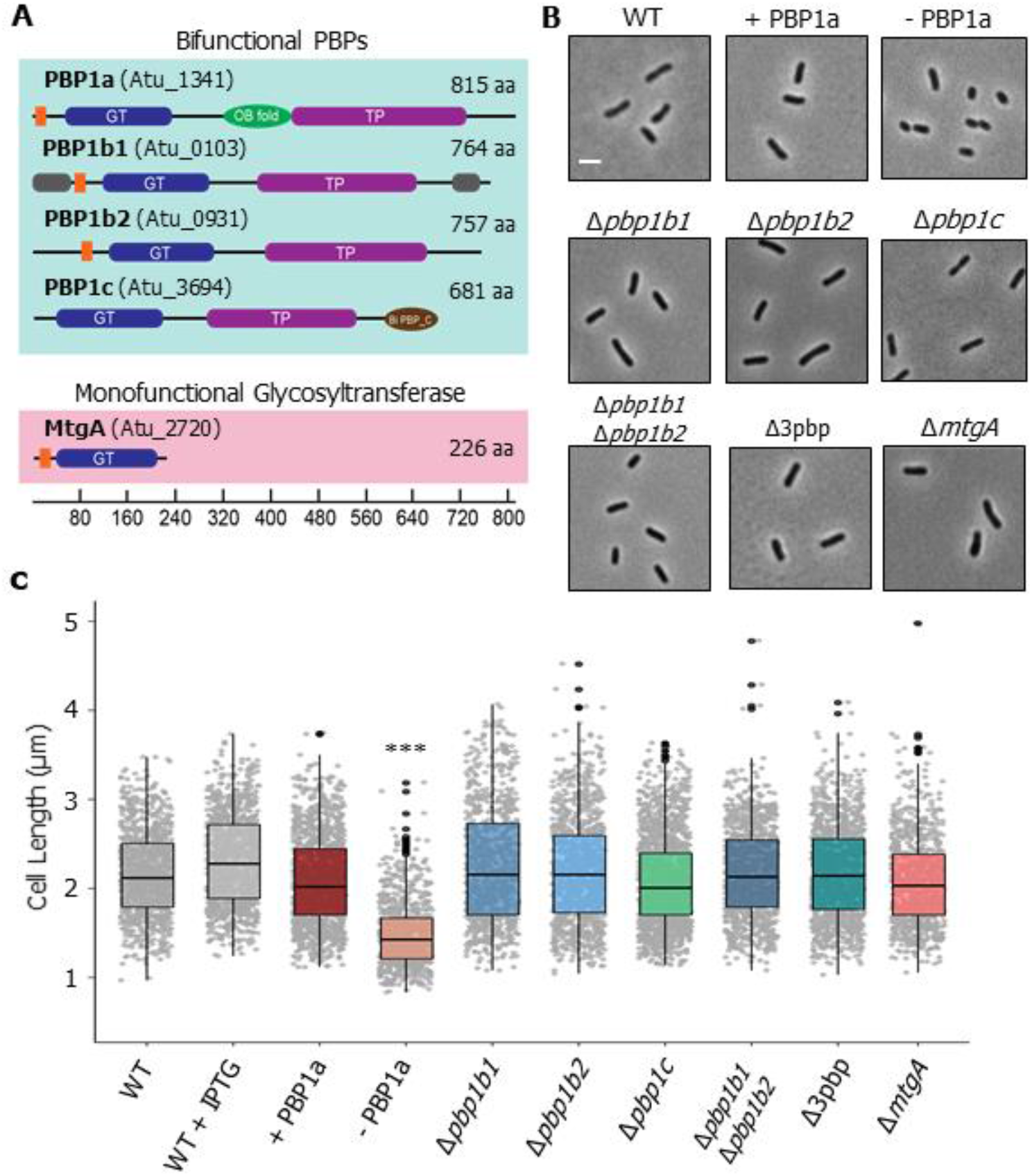
Functional characterization of PG synthases in *A. tumefaciens*. (**A**) Domain structure of the putative PG biosynthesis enzymes showing the transmembrane (orange), glycosyltransferase (GT, PF00912), transpeptidase (TP, PF00905), OB-like (PF17092), and biPBP_C (PF06832) domains. The regions of intrinsic disorder (grey) as predicted by MoBiDB are also shown [23]. The scale indicates the length in amino acids (aa). The corresponding ATU numbers are listed in parentheses beside each gene name. (**B**) Phase microscopy images showing the phenotypes of PG synthase mutants. Each strain was grown to exponential phase, spotted on an ATGN agar pad (ATGN is a minimal medium with glucose and (NH4)2SO4), and imaged by phase microscopy. Scale bar: 2 μm. (**C**) Cell length distributions of PG synthase mutants. The indicated strains were grown as in B and subjected to cell lengths measurements using MicrobeJ [24]. The data are represented as box and whisker plots in the style of Tukey [25], and visualizes five summary statistics (the center line is the median, the two hinges correspond to the first and third quartiles (the 25^th^ and 75^th^ percentiles), the two whiskers (representing the smallest and largest values no further than 1.5 times the interquartile range), and all “outlying” points are plotted individually. The PBP1a depletion strain grown in the presence of IPTG is referred to as + PBP1a and the depletion strain grown in the absence of IPTG is referred to as - PBP1a. Distributions of cells significantly different from wildtype (WT) are indicated (***; One-Way ANOVA with Bonferroni correction, p > 2*10^16). n = > 800 cells per strain.

**Figure 2.**
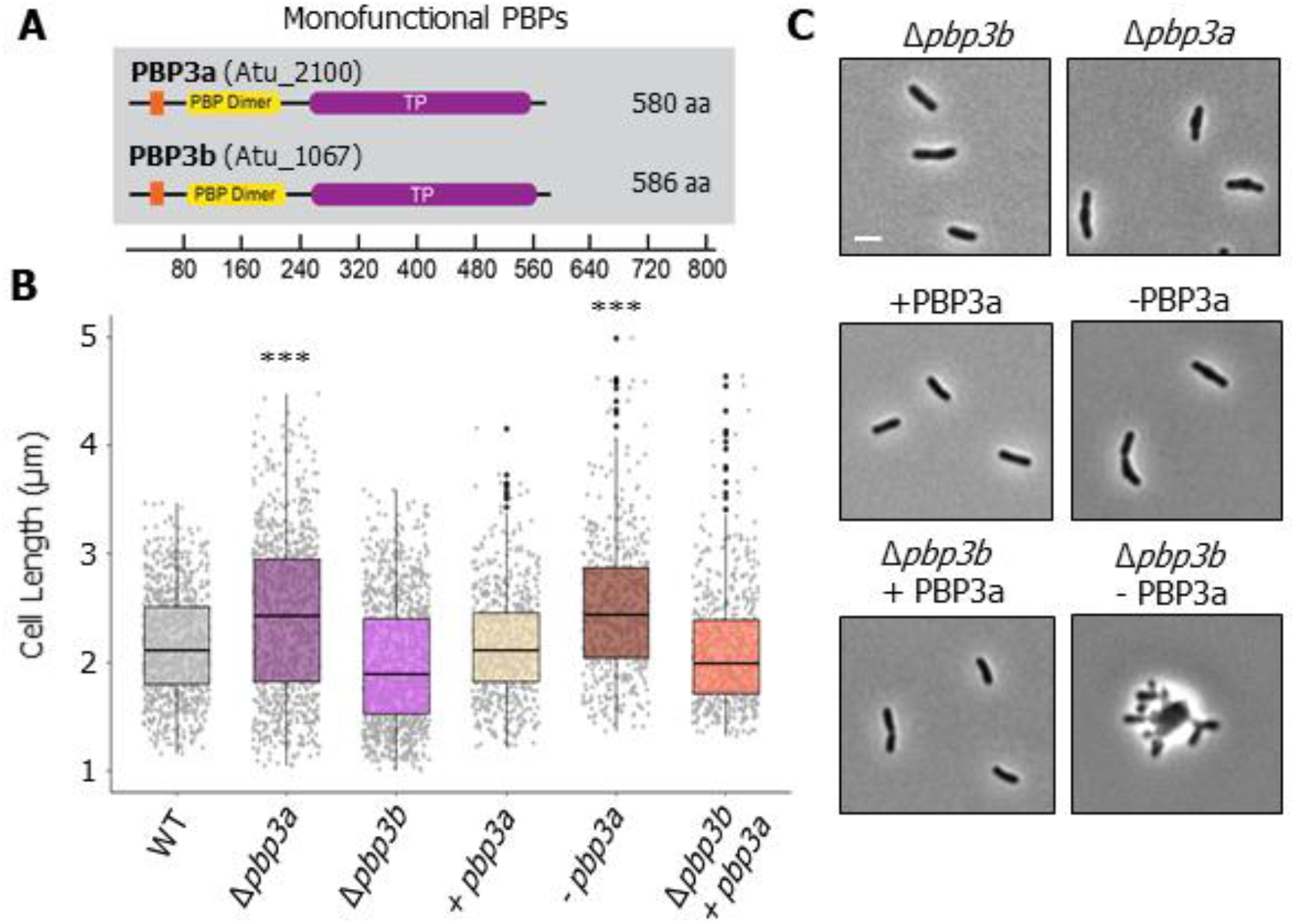
Functional characterization of monofunctional synthases PBP3a and PBP3b. (**A**) Domain structure of the putative PG biosynthesis enzymes showing the transmembrane (orange), transpeptidase (TP, PF00905), and PBP dimer (PF03717) domain. The scale indicates the length in amino acids (aa). The corresponding ATU numbers are listed in parentheses beside each gene name. (**B**) Phase microscopy images showing the phenotypes of PG synthase mutants. Each strain was grown to exponential phase, spotted on an ATGN agar pad, and imaged by phase microscopy. Scale bar: 2 μm. (**C**) Cell length distributions of PG synthase mutants. The indicated strains were grown to exponential phase, spotted on an agar pad, imaged by phase microscopy, and subjected to cell lengths measurements using MicrobeJ [24]. The data are shown as box and whisker plots in the style of Tukey [25]. Distributions of cells significantly different from wildtype (WT) are indicated (***; One-Way ANOVA with Bonferroni correction, p< 2*10^16). n = > 500 per strain.

Consistent with a previous transposon screen [26] it was not possible to obtain a PBP1a deletion. We therefore constructed a PBP1a depletion strain by introducing a copy of the PBP1a-encoding gene, under an IPTG-inducible promoter, at a heterologous site on the chromosome and subsequently succeeded in creating an in-frame deletion of the native gene encoding PBP1a in the presence of the IPTG inducer [27]. The PBP1a depletion strain grown in the presence of IPTG is referred to as + PBP1a and the depletion strain grown in the absence of IPTG is referred to as - PBP1a. We confirmed depletion of PBP1a in the absence of IPTG using Bocillin-FL, a fluorescent penicillin derivative. Two bands were observed that could correspond to the predicted molecular weight of PBP1a (~88 kDa), but only the second band was absent in PBP1a-depleted cells and likely represents PBP1a protein (Supplementary Figure 1A). Strikingly, cells depleted of PBP1a for 16 hours lost their rod shape, becoming shorter (Figure 1A, B) and wider (Supplementary Figure 1B) and had a severe viability defect, as measured by spotting serial dilutions, compared to the same strain when PBP1a is induced (Supplementary Figure 1C). The addition of IPTG to wild-type *A. tumefaciens* led to a slight increase in the median cell length (Figure 1B, 1C) compared to wildtype alone but had no effect on cell viability or rod shape (Supplementary Figure 1B, 1C).

Of the remaining bifunctional PBPs, single deletions of the genes encoding PBP1b1, PBP1b2, and PBP1c had no effect on cell length (Figure 1B, C) or cell viability (Supplementary Figure 1D). Similarly, a double mutant of PBP1b1 and PBP1b2 or a triple mutant of PBP1b1, PBP1b2 and PBP1c (referred to as Δ3pbp) had no obvious mutant phenotype with respect to cell length (Figure 1B, C) or cell viability (Supplementary Figure 1D). Thus, despite all of the predicted bifunctional PBPs being produced during exponential growth of *A. tumefaciens* (Supplementary Figure 1E), our data indicate that only PBP1a makes a major contribution to cell growth under standard laboratory conditions. Similarly, deletion of the monofunctional GTase encoding *mtgA* produced cells of normal length (Figure 1B, C). The lack of a readily observed phenotype in the *ΔmtgA* strain is consistent with findings in *E. coli* and *Hyphomonas neptunium* [28–30]. Together, these data suggest that the bifunctional enzyme PBP1a, which likely has both GTase and TPase activities, fulfils the role of RodA and PBP2, as the primary PG synthase required for polar elongation in *A. tumefaciens*.

### Class B synthases PBP3a and PBP3b are required for cell division

Incorporation of PG at the septum prior to cell division typically requires synthesis enzymes that are distinct from the cell elongation machinery. Although *A. tumefaciens* lacks the cognate SEDS-PBP pair that is typically required for elongation, the SEDS protein FtsW and PBP3, which are required for cell division, are conserved. While most bacteria possess a single, essential *ftsI* gene that encodes PBP3, some Rhizobiales, including *A. tumefaciens*, encode two FtsI homologs (PBP3a and PBP3b) (Figure 2A). *pbp3a* resides in the *mra* operon of cell division and cell envelope biogenesis genes similar to most *ftsI-encoding* homologs [31], while PBP3b is encoded as a monocistronic gene elsewhere in the genome. This raises the possibility that the second monofunctional transpeptidase (PBP3b) may serve the role of a PBP2 homolog that functions in polar elongation. Thus, we sought to determine if either of the two class B PBP homologs were required for cell division or polar elongation in *A. tumefaciens*. Saturating transposon mutagenesis in LB medium indicated that PBP3a was likely essential [26]; however, it was possible to make a deletion of *pbp3a* when cells were grown in minimal medium. These observations suggest that the PBP3a-encoding gene was conditionally essential.

In minimal medium, deletion of *pbp3a* caused a severe cell viability defect (Supplementary Figure 1F). In addition, cells were longer (Figure 2B, C), and were frequently observed to adopt branching or bulging morphologies; these features are a hallmark of cell division defects in *A. tumefaciens* [32, 33]. In contrast, deleting *pbp3b* had no effect on cell viability (Supplementary Figure 1F) or cell length (Figure 2B, C). Together, these results indicate that PBP3a is the major class B PBP contributing to cell division in *A. tumefaciens*. Since the deletion of the gene encoding PBP3a did not fully inhibit cell division, we hypothesized that PBP3b may be able to partially compensate for the loss of PBP3a. To address this possibility, we first created a PBP3a depletion strain; when this strain was grown in the absence of IPTG for 24 hours, it phenocopied the cell viability and cell length defects of the *pbp3a* deletion mutant (Figure 2B, C). We then depleted PBP3a in a *Δpbp3b* mutant background, and found that cells not only failed to divide, but also swelled at the mid-cell before lysing (Figure 2C, Supplemental movie 1), indicating that PBP3a and PBP3b both contribute to septal PG biosynthesis during cell division. The phenotype observed in the absence of both PBP3a and PBP3b was remarkably similar to the phenotype observed during depletion of FtsW in *A. tumefaciens* [32]. FtsW possesses GTase activity, and is a major synthase required for cell division [12, 34]. Consistent with current models of cell division, we hypothesize that FtsW provides the GTase activity while PBP3a and PBP3b provide the dd-transpeptidase (TPase) activity necessary for proper septal PG biosynthesis during cell division in *A. tumefaciens*. In all, our findings support a model in which PBP3a can sustain proper cell division in the absence of PBP3b, and that, while PBP3b contributes to septal PG synthesis, it cannot fully compensate for the loss of PBP3a. Thus, both class B PBPs contribute primarily to septal PG biosynthesis.

### Development of new fluorescent cell wall probes to monitor PBP activity

Traditional fluorescent d-amino acid (FDAA) probes are an exceptionally useful tool for investigating the patterning of cell wall synthesis in diverse microbes [35]. However, FDAAs report on the activity of extracellular/periplasmic dd and ld transpeptidases, and as a result can be incorporated into the PG in a growth-independent mechanism [6]. Here, we have developed fluorescent d-amino acid dipeptide (FDAADs) probes to observe nascent sites of PG crosslinking in living cells, thus eliminating the need for click chemistry that is required when using traditional d-amino acid dipeptide probes (DAADs) [6, 36]. DAADs are incorporated into the cell wall precursors by the cytoplasmic MurF ligase and probe incorporation reports specifically on nascent PG synthesis [36–38]. The resulting lipid II-linked modified precursor is most likely covalently crosslinked to an existing glycan strand through the activity of bifunctional PBPs (Figure 3A). The FDAAD probe HADA—DA successfully labeled the PG of several bacteria, including *B. subtilis, E. coli, Streptomyces venezuelae and A. tumefaciens* (Figure 3B, C), and we demonstrated and evaluated PG labeling with four additional FDAADs of different sizes and molecular weights in diverse species (Supplemental figure 2A-D). In addition, *S. venezuelae*, a polar growing Actinobacteria, was short pulsed with three different FDAADs to illustrate that these probes report on the newest PG synthesis activity (Figure 3B), similar to FDAAs [39, 40]. Given their cytoplasmic mechanism of incorporation, complementary to FDAAs (which are incorporated by transpeptidases periplasmically) [6], these probes will be particularly useful to distinguish between growth-dependent and growth-independent PG crosslinking in species with a higher proportion of extracellular LDT activity, such as polar-growing bacterial species.

**Figure 3.**
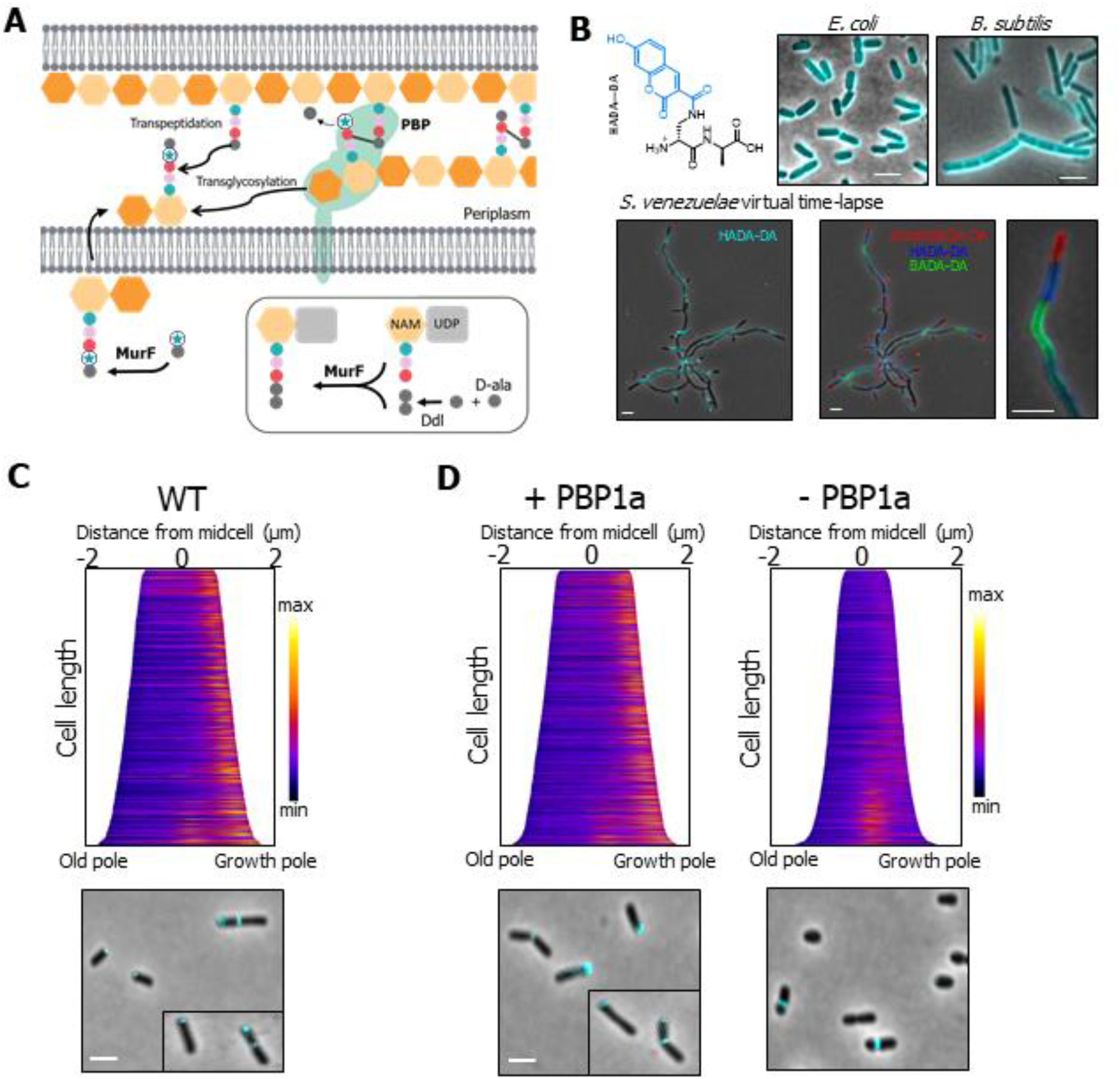
Fluorescent d-amino acid dipeptide (FDAAD) labeling is absent from the growth pole in the PBP1 depletion. (**A**) Schematic of the incorporation pathway of **f**luorescent **d**-**a**mino **a**cid **d**ipeptides (FDAADs), which are incorporated into the muropeptide precursor molecule in the cytoplasm by MurF ligase. The modified muropeptide precursor is flipped across the membrane and the activity of a bifunctional PBP crosslinks the new PG monomer into the existing PG sacculus. (**B**) Top row: structure of the FDAAD HADA—DA and merged phase and fluorescent channels of labeling patterns of HADA—DA in *E.coli* and *B. subtilis*. Bottom row: short pulse labeling of *S. venezuelae* sequentially labeled first with BADA—DA, followed by HADA—DA and Atto610DA—DA. (**C**) Demographs of wild-type (WT) cells depict incorporation of FDAADs at a population level. Median profiles of the fluorescence channel are stacked and ordered by cell length n = 513. (**D**) Demographs depict incorporation of FDAAD in the PBP1a depletion strain grown in the presence or absence of IPTG labeled + PBP1a and - PBP1a, respectively. Merged phase and fluorescent channels of cells with representative polar and septum labeling of FDAAD are shown to the right of each demograph. Scale bars: 2 μm. n = > 1000 per strain

### PBP1a is the major synthase incorporating PG at the pole

*A. tumefaciens* was labeled for ~5% of the cell cycle with HADA—DA. As expected, cells exhibit labeling at the growth pole during elongation, and at the septum in cells undergoing cell division (Figure 3C). This pole-to-septum labeling pattern is consistent with other cell wall labeling methods including fluorescent d-amino acids (FDAAs) [41] and d-cysteine labeling [1]. Similar to wild-type cells, a pole-to-septum labeling pattern was observed in the Δ3pbp mutant, consistent with a limited role for these class A PBPs in polar growth under the conditions tested (Supplementary Figure 3).

In stark contrast to wild-type (Figure 3C) or PBP1a replete cells (Figure 3D), pole-specific labeling with HADA—DA was almost completely absent from cells depleted of PBP1a for 20 hours (Figure 3D). Similar results were observed with NADA—DA and BADA—DA (Supplemental Figure 4A, B). Consistent with these observations, we tracked the growth of PBP1a-depleted cells for seven generations using a microfluidic device and found that the reduction in cell length occurred first for the new pole daughter cell and likely resulted from the loss of polar PG insertion by PBP1a (Supplemental Figure 3A, Supplemental movie 2). We thus concluded that PBP1a is the major PG synthase required for polar PG incorporation. Notably, PBP1a-depleted cells have more robust FDAAD labeling at the septum (Figure 3D), suggesting that additional glycosyltransferase enzymes remain active at the site of cell division.

### ld-transpeptidases contribute to growth-independent and growth-dependent PG modification

In contrast to FDAAD incorporation, which is linked to the PG precursors in the cytoplasm, the more conventional fluorescent d-amino acid (FDAA) labeling occurs in the periplasm through either dd-transpeptidase reactions carried out by PBPs or ld-transpeptidase reactions carried out by LDTs [41, 42]. As expected, wild-type and Δ3pbp cells labeled with a short pulse of the FDAA probe HADA show the characteristic pole-to-septum labeling pattern that is typical of *A. tumefaciens* growth (Figure 4A, Supplementary Figure 3). Notably, FDAADs label discrete regions at the pole and midcell, while FDAA labeling is more prominent in the sidewalls, particularly of the new pole compartment prior to cell division, while the old pole remains unlabeled. (Figure 4A white arrow). This labeling pattern is consistent with the reported growth-independent incorporation of FDAAs by LDTs [6]. Interestingly, we observed that a majority of growth-independent labeling occurred preferentially in the new pole compartment. To further support this idea, we labeled *A. tumefaciens* for 60 minutes with either HADA or HADA—DA and compared the labelling pattens (Supplemental figure 5). After a 60-minute incubation, HADA—DA labeling was primarily found in distinct regions at the pole and midcell, with little sidewall labeling, similar to a short pulse and consistent with areas of new synthesis. In contrast, the HADA labelling was much brighter, and fully labeled the sidewalls. HADA labeling was particularly enriched along the sidewalls of the new-pole compartment. Our observations during short and long-pulse labeling experiments suggests that, in addition to growth-dependent pole and midcell labeling, LDTs contribute to spatially distinct crosslinking of the cell wall, particularly along the sidewalls of the new pole daughter cell.

**Figure 4.**
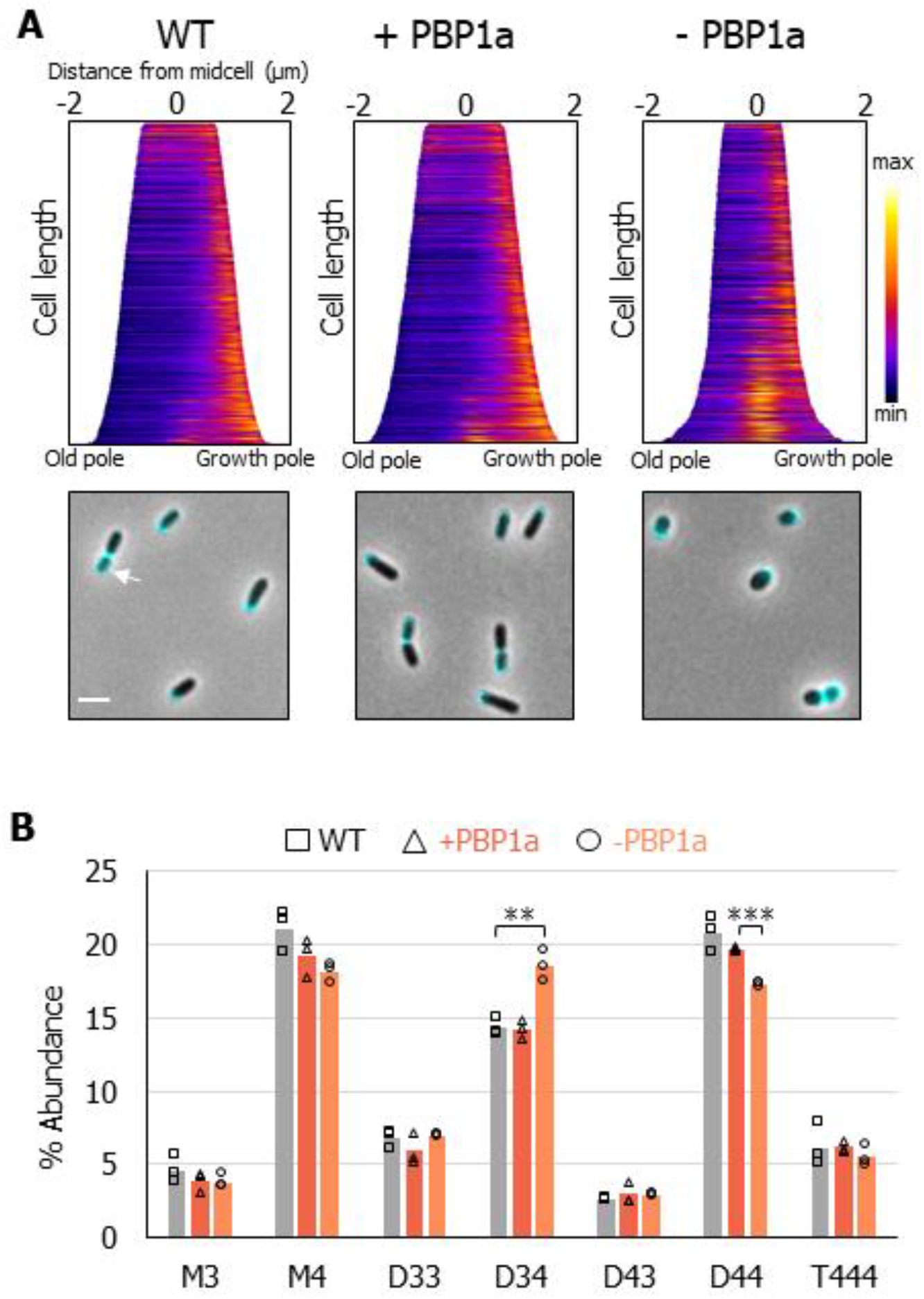
Fluorescent d-amino acid (FDAA) and PG composition analysis illustrates a role for LDTs in polar growth. (**A**) Demographs of wildtype (WT) and the PBP1a depletion strain grown in the presence or absence of IPTG labeled + PBP1a and - PBP1a, respectively. Demographs depict incorporation of FDAA at a population level. Median profiles of the fluorescence channel are stacked and ordered by cell length wildtype n = > 600 per strain. Merged phase and fluorescent channels of cells with representative polar and septum labeling by FDAAs are shown below each demograph. Scale bar: 2 μm. (**B**) Bar graphs depicting the average abundance of muropeptides obtained by UPLC analysis from wildtype, and the PBP1a depletion strain grown in the presence or absence of IPTG for 16 hours. Major muropeptides are labeled M, monomers; D, dimers; and T, trimers. Numbers indicate the length of the muropeptide stems and the position of cross-links in dimers and trimers. Data shown are averages taken from analysis of three independent biological samples. Samples that are statistically significant are indicated (One-Way ANOVA with Tukey’s multiple comparison test, ** p<0.005, *** p<0.0005). p-value between + PBP1a and - PBP1a for D34 was 0.0588 and between WT and - PBP1a for D44 was 0.072.

We next sought to test whether FDAA labeling at the growth pole was absent in the PBP1a depletion strain. In contrast to the absence of polar incorporation that was seen following FDAAD labeling, cells depleted of PBP1a labeled robustly at the growth pole with HADA (Figure 4A). Since PBP-mediated incorporation of FDAADs is absent from the growth pole in the PBP1a depletion (Figure 3D, Supplemental Figure 4A, B), the major enzymes incorporating HADA at the pole are most likely LDTs, consistent with recent findings for incorporation of FDAAs in *E. coli* [42]. Additionally, the activity of LDTs has been shown to be functionally linked to the activity of class A PBPs [42, 43]. Therefore, our findings indicate that ld-transpeptidation contributes to PG crosslinking at the growth pole.

### Depletion of PBP1a or loss of PBP1b2 leads to increased levels of ld-crosslinks in PG

Since depletion of PBP1a led to dramatic reduction in the polar incorporation of FDAADs, while maintaining polar FDAA incorporation we hypothesized that a decrease in the activity of PBPs relative to LDTs may lead to altered PG composition. We grew the PBP1a depletion strain in the presence or absence of IPTG for 16 hours, collected the cell wall fraction, and analyzed muropeptides by ultra-performance liquid chromatography (UPLC). The major muropeptides found in wild-type *A. tumefaciens* PG, included monomeric (M), dimeric (D) and trimeric (T) muropeptides (Figure 4B). PG from PBP1a-depleted cells had significantly reduced levels of muropeptides with dd-crosslinks, as seen by the ~3% reduction in D44 abundance (Figure 4B). As expected, this observation confirmed that depleting PBP1a leads to decreased dd-transpeptidase activity. Depletion of PBP1a also resulted in an increase in muropeptides containing ld-crosslinks, as indicated by the ~4% increase in D34 abundance (Figure 4B). A similar decrease in D44 abundance and increase in D34 abundance was also observed in the *Δpbp1b2* (Supplemental Figure 4A), but not in the single deletions of *pbp1b1, pbp1c, pbp3a* or *pbp3b* (Supplemental Figure 4B). Therefore, the compositional changes in the double and triple mutant strain could likely be attributed to the deletion of *pbp1b2*. These results implicate PBP1b2 as an important dd-transpeptidase enzyme in *A. tumefaciens* and suggest that increased LDT activity may be a general response to decreased dd-transpeptidase levels, and not necessarily a specific response to depletion of PBP1a. Furthermore, since we saw an increase in muropeptide D34 but not D33 (Figure 4B), we hypothesize that decreased dd-transpeptidase levels activate only a subset of LDTs, and that another group of LDTs may function along with PBP1a during polar growth, consistent with the observation of HADA labeling at the growth pole.

### Faropenem treatment inhibits polar growth in the Rhizobiales

Since species in the Rhizobiales have a large number of ld-transpeptidase enzymes (*A. tumefaceins* encodes 14 LDTs), deleting them all is a significant undertaking. β-lactam antibiotics are one of the most widely used classes of antibiotics that target cell wall synthesis enzymes, and these have been primarily studied for their ability to target PBPs [44]. A subclass of β-lactam antibiotics known as the carbapenems, including the penem antibiotic faropenem, are, however, useful for probing the activity of LDTs [45, 46]. Therefore, to characterize the global contribution of LDT activity during *A. tumefaciens* growth, we investigated the effect of five carbapenem antibiotics and one penem antibiotic on cell growth and morphology. In *A. tumefaciens*, we find that treatment with meropenem or faropenem leads to an overall decrease in PG crosslinkage, including both ld- and dd-crosslinks (Figure 5A, Supplemental Figure 5), suggesting that the targets of these drugs impact cell wall biosynthesis.

**Figure 5.**
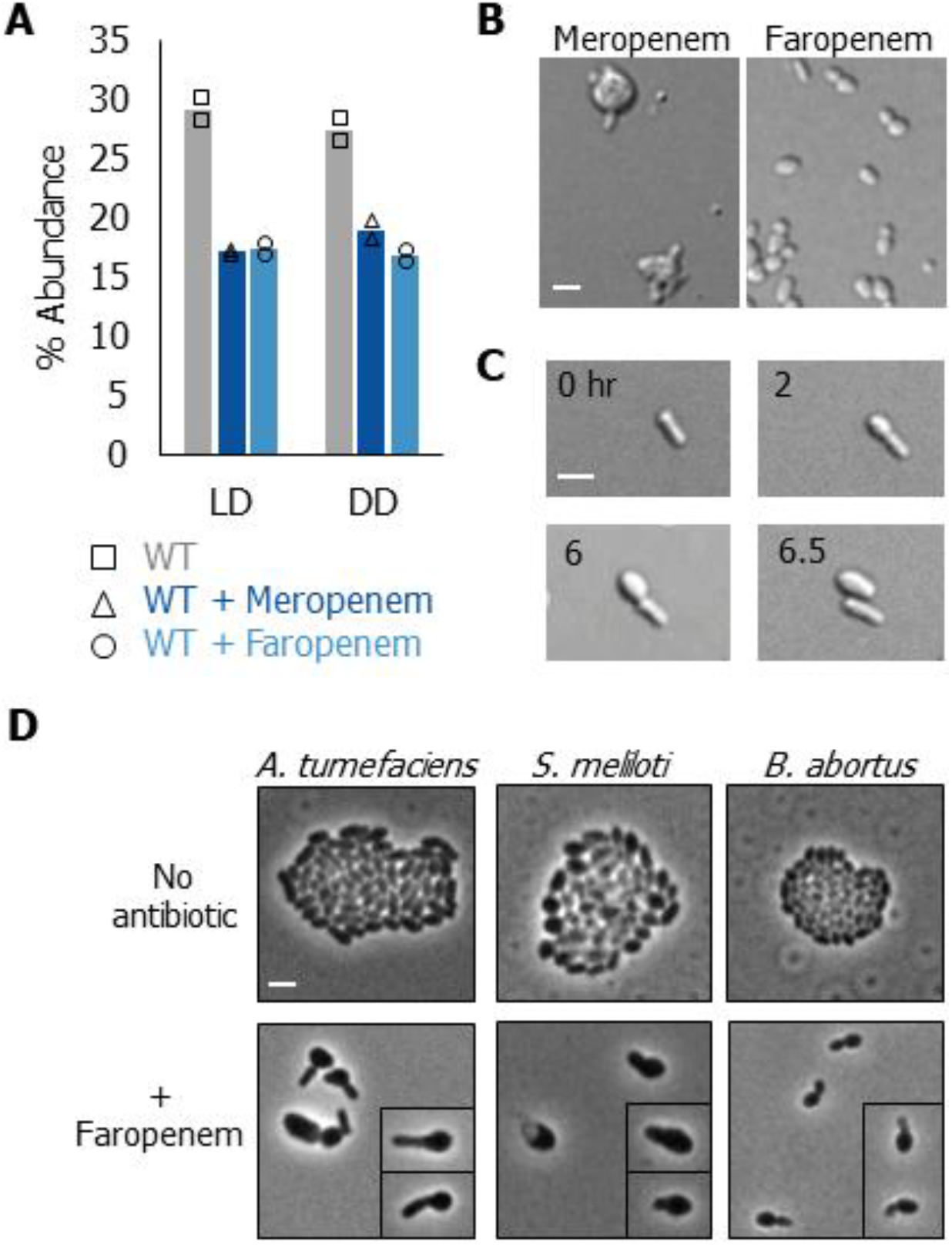
Phenotypic characterization of carbapenem and penem antibiotic treatments. (**A**) Abundance of total ld- and dd-crosslinkages in PG isolated from wild-type cells, wild-type cells grown in the presence of 1.5 μg/mL meropenem for 4 hours, and wild-type cells grown in the presence of 1.5 μg/mL of faropenem for 6 hours. Timepoints were chosen based on the onset of phenotypic changes. Data shown are the total abundance of the muropeptides containing ld- or dd-crosslinks from analysis of two independent samples. (**B**) Representative images of wild-type cells grown in the presence of 1.5 μg/mL meropenem and faropenem. Cells were incubated with antibiotics for 24 hours, then spotted on a 1% agarose pad and imaged using DIC microscopy. Scale bar: 2 μm (**C**) Time-lapse microscopy of wild-type cells spotted on a 1% ATGN agarose pad supplemented with 1.5 μg/mL of faropenem; images were acquired every 10 minutes. Indicated time in hours is shown. Scale bar: 2 μm (**D**) Representative phase microscopy images of *A. tumefaciens, S. meliloti*, or *B. abortus* grown overnight in ATGN, Tryptone-Yeast (TY) Extract or Brucella Broth, respectively, to an OD_600_ of 0.6 and spotted on a 1% ATGN agarose pad with or without 1.5 μg/mL faropenem. Cells were imaged after 16 hours of growth. Scale bar: 2 μm.

To better understand the impacts of these antibiotics, we observed morphological changes induced by drug treatment. Treatment with sub-minimum inhibitory concentrations (MIC) treatments with any of five carbapenem antibiotics: meropenem, imipenem, doripenem, ertapenem, or tebipenem for 24 hours induced mid-cell swelling (Figure 5B, Supplementary Figure 6). These data indicated that these carbapenem antibiotics may target an enzyme(s) with a specific role at the septum during cell division. Interestingly, faropenem-treated cells became wider and rounder after 24-hour exposure (Figure 5A), which pointed to the cellular target of faropenem as being important for the maintenance of rod shape during polar growth. In agreement, time-lapse microscopy of faropenem-treated cells revealed a loss of rod shape preferentially in the daughter compartment. Remarkably, following cell division the daughter cell which inherits the growth pole is large and round whereas the daughter cell inheriting the old growth pole retains its rod shape (Figure 5B, Supplemental movie 3). This phenotype was distinct from that associated with the treatment of the other carbapenem antibiotics and indicated that the cellular target(s) of faropenem was important to maintain proper PG integrity in the growth pole compartment.

The phenotype of faropenem-treated cells, along with the growth-independent labelling of HADA along the sidewalls of the new pole (Supplemental figure 6) suggested that the old and new cell compartments may have a distinct repertoire of PG synthesis enzymes that help to confer polar identity between the daughter cells. To determine if the cellular target(s) of faropenem were conserved in other Rhizobiales we treated the closely related plant symbiont *S. meliloti* and the obligate intracellular pathogen *B. abortus* with sub-lethal concentrations of faropenem. We observed swelling of the growth pole in *A. tumefaciens, S. meliloti*, and *B. abortus* (Figure 5D). Thus, we have identified the β-lactam antibiotic faropenem as a specific antibiotic inhibitor of polar growth among the Rhizobiales. Altogether, these data suggest that cell wall enzymes that are important for polar growth have a conserved role in agriculturally and medially important species of Rhizobiales.

### The essentiality of PBP1a is conserved among the Rhizobiales

Since faropenem targeting of the growth pole machinery is conserved, we sought to determine if the essential role of PBP1a is also conserved in other Rhizobiales. We found that treatment of *B. abortus* with the PBP1a-specific GTase inhibitor flavomycin (moenomycin) [47] led to growth arrest and the formation of large, round cells (Figure 6B). Consistent with our findings that treatment of *B. abortus* with the PBP1a inhibitor flavomycin causes cells to lose their rod-shape, a transposon mutagenesis screen of *B. abortus* predicted that out of the bifunctional PBPs, only PBP1a may be essential for growth [48]. In line with this, Bandara and colleagues were unable to obtain a PBP1a mutant in *Brucella melitensis*, indicating that PBP1a is likely essential for viability [49].

**Figure 6.**
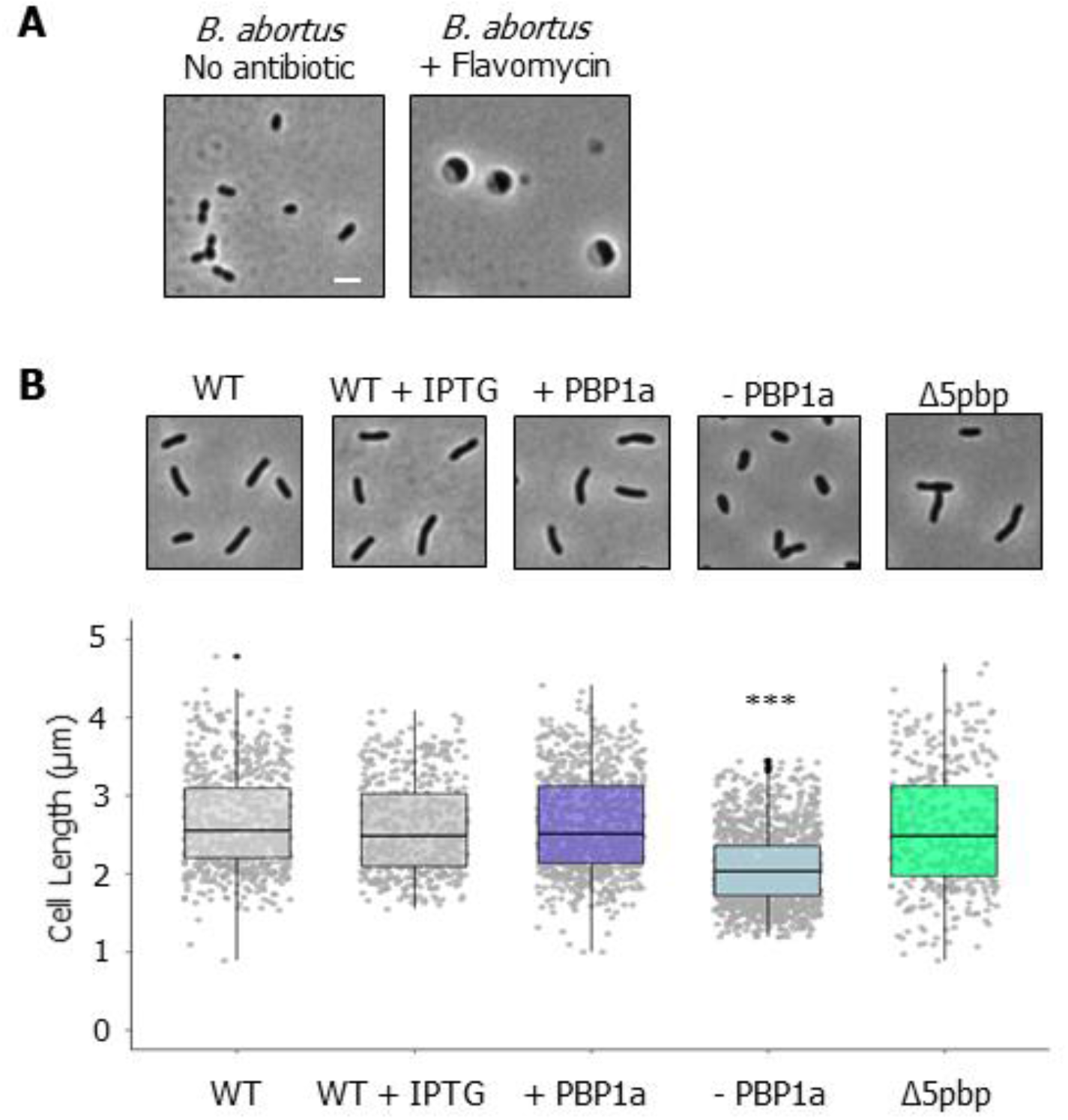
Mechanisms of polar growth are conserved in the Rhizobiales *Brucella* and *Sinorhizobium*. (**A**) Representative phase microscopy images of *B. abortus* grown overnight without antibiotic, with 20 μg/mL of flavomycin, or with 1.5 μg/mL of faropenem, after 16 hours cells were spotted on a 1% Brucella Broth agarose pad and imaged. (**B**) Top panel: phase microscopy images showing the phenotypes of *S. meliloti* wildtype (WT) and PBP mutants. Each strain was grown to exponential phase, spotted on a 1% TY agarose pad, and imaged by phase microscopy. Bottom panel: cell length distributions of PG synthase mutants. The indicated strains were grown to exponential phase, spotted on an agarose pad, imaged by phase microscopy, and subjected to cell lengths measurements using MicrobeJ [24]. The data are shown as box plots in the style of Tukey [25]. Distributions of cells significantly different from wildtype are indicated (***; One-Way ANOVA with Bonferroni correction, p< 2*10^16). n = >400 per strain.

Next, we explored the function of PBPs in the closely related plant symbiont *S. meliloti. S. meliloti* encodes six bifunctional PBP homologs, including one PBP1a homolog, four PBP1b homologs, and one PBP1c homolog. We constructed a Δ5pbp mutant, which is lacking all four PBP1b homologs and the PBP1c homolog. The Δ5pbp mutant had a median cell length similar to that of wild-type *S. meliloti*, but with a slightly broader distribution of cell lengths, with ~88% of cells falling between 1.5 and 4 μm compared to ~97% of WT cells falling between these cell lengths. Interestingly, the Δ5pbp mutant retained its rod shape, similar to the Δ3pbp mutant of *A. tumefaciens*, suggesting that these enzymes contribute minimally to sustaining proper rod shape under standard growth conditions.

Similar to our findings for *A. tumefaciens*, we were unable to make a deletion of *pbp1a*, so we constructed a PBP1a depletion strain. Depletion of PBP1a led to a severe viability defect (Supplementary Figure 7A) and cells became shorter (Figure 6B) and wider (Supplementary Figure 7B). Thus, PBP1a is essential for polar growth and maintenance of proper rod shape in both *S. meliloti* and *A. tumefaciens*. Taken together, our data from *B. abortus, S. meliloti* and *A. tumefaciens* support the notion that PBP1a plays an essential and conserved role in polar growth among the Rhizobiales.

## DISCUSSION

Bacteria employ widely diverse growth strategies. Unipolar growth, or incorporation of new cell wall material at a single pole, is a shared mode of growth among the Rhizobiales, but the mechanisms that drive polar PG insertion remain poorly understood. Most bacteria have multiple class A PBPs with semi-redundant functions and growth is supported by the presence of any one of the PBPs [50] In *C. crescentus*, four class A PBPs can support growth when expressed alone. [51] In *B. subtilis*, all 4 class A PBPs are dispensable and the SEDS protein RodA has likely taken over the essential function of the class A PBPs during elongation [52] In contrast, here we identified the class A PBP1a homolog as an essential protein needed to synthesize PG at the growth pole in *A. tumefaciens* and related genera, despite the presence of other class A PBPs. Current findings support the proposal that in laterally growing bacterial species, PBP1a homologs act as autonomous entities involved in PG remodeling or repair and do not function as a part of the core elongasome [15, 16]. Instead, the monofunctional glycosyltransferase RodA and the monofunctional transpeptidase PBP2 are required to maintain rod-shape [10, 53]. In contrast, we find that in the Rhizobiales, which lack MreB, RodA and PBP2 homologs, the bifunctional enzyme PBP1a is an essential core component of the elongasome of polar growing Rhizobiales. Loss of class A PBPs in *E. coli* or *B. subtilis* led to a decrease in cell width [54, 55]. Conversely, depletion of PBP1a in *A. tumefaciens* and *S. meliloti* led to a significant decrease in cell length and an increase in cell width (Figure 1B, C, Supplementary Figure 1B). This indicates that cells lacking PBP1a have a shorter period of cell elongation, and as a result, likely spend more time synthesizing the septum (Figure 3C, Supplementary Figure 3A, B). Thus, the regulation of PG synthesis that governs cell length and cell width in the Rhizobiales utilizes a novel mechanism compared to well-studied model bacteria. Perhaps because class A PBPs function independently of cytoskeletal complexes [10], PBP1a was freely available to assume the role of the primary enzyme driving polar PG synthesis prior to the loss of the *mre* operon. Notably, the other bifunctional PBPs minimally contributed to the maintenance of proper rod shape under the conditions tested in *A. tumefaciens* and *S. meliloti*. However, it is likely that the additional class A PBPs may make dedicated contributions under specific growth conditions. For example, in *E. coli*, PBP1a homologs are required for optimal growth in alkaline pH, while PBP1b homologs are required under acidic conditions [56]. Since the plant rhizosphere is an acidic environment [57], it is possible that the remaining bifunctional PBPs have specialized functions to maintain growth when bacteria are associated with a plant host. Additionally, duplication of the monofunctional PBP3 homolog among the Rhizobiales is restricted to a few species that interact with plants, suggesting this is not a broad solution to the loss of PBP2, but that PBP3b homologs may contribute more significantly to bacterial growth in plantae. In agreement with this idea, we found that of the two monofunctional PBP homologs (PBP3a and PBP3b), PBP3a - a clear homolog of the division-specific PBP3 - contributes to cell division, while deletion of *pbp3b* plays a minimal role in cell growth or cell shape under standard laboratory conditions but forms a synthetic lethal pair with PBP3a that functions in cell division.

In laterally growing bacterial species, the role of scaffolding PG synthase enzymes during elongation is fulfilled in part by the actin homolog MreB [58]. Homologs of MreBCD are absent in the Rhizobiales; thus, how PBP1a is recruited to the growth pole and how its activity is regulated remains unexplored. Recently, GPR (for Growth Pole Ring), a large (~226 kDa) apolipoprotein with similarity to the polar organizing protein TipN from *C. crescentus*, was reported to form a ring at the growth pole in *A. tumefaciens*, with depletion of this protein leading to rounded cells [59]. This phenotype implicates GPR as a possible candidate to scaffold PG enzymes during elongation. In addition, PG synthesis by PBP1a also requires hydrolysis of the existing sacculus to allow for insertion of new muropeptides. A dd-endopeptidase (RgsM) that is predicted to have hydrolysis activity was recently shown to be essential for polar growth in *S. meliloti* [60], and thus represents an interesting candidate for polar PG hydrolysis.

Several lines of evidence suggest that FDAAs are primarily incorporated into the cell wall through remodeling by LDTs [41, 42, 61, 62]. Cells depleted of PBP1a label robustly at the growth pole with FDAAs, independently of PBP-mediated FDAAD labeling. We also found that FDAAs label WT cells along the sidewalls of the new pole much more brightly than the old pole. This points to a role for LDTs in not only polar growth, but also sidewall remodeling of the new pole daughter cell (Figure 7).

**Figure 7.**
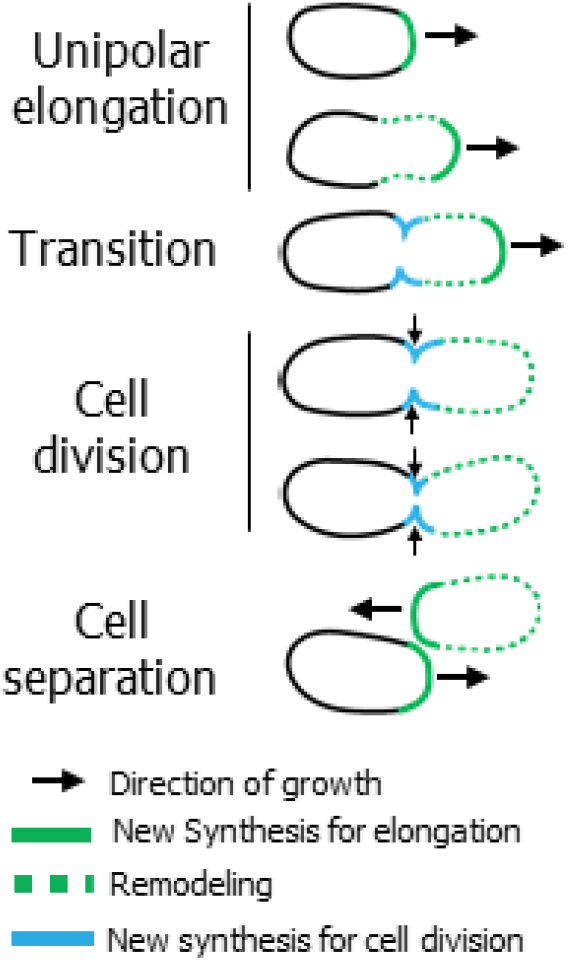
Model of cell wall synthesis in the Rhizobiales. **Unipolar elongation:** *A. tumefaceins* elongates from a single pole using the action of an essential PBP1a homolog. As the cell gets longer, the sidewalls of the new pole begin to be remodeled in a growth-independent manner, through the activity of ld-transpeptidases. **Transition:** prior to cell division growth at the pole is terminated and growth at the midcell is initiated, while remodeling of the new pole compartment continues. **Cell division:** during cell division new PG is added at the midcell through the transpeptidase activity of two class B PBPs (with PBP3a being the primary synthase) and the glycosyltransferase FtsW. **Cell separation:** after cell separation, the new poles resume polar elongation from what was the site of cell division. Continuous remodeling of the new pole daughter cell occurs throughout the cell cycle and is inherited into the new pole daughter cell.

Collectively, our results indicate that at the pole in *A. tumefaciens* FDAAD probes report on nascent PG synthesis by PBP1a, whereas FDAAs are incorporated by the localized activity of LD-transpeptidases on the bacterial cell surface. These findings demonstrate the usefulness of FDAAD probes for distinguishing between growth-dependent, nascent cell wall synthesis and growth-independent cell wall remodeling. In particular, the growth pattern of species for which a high proportion of growth-independent remodeling occurs can be obscured by FDAAs. Therefore, FDAADs are more useful in distinguishing nascent PG synthesis mediated by PBPs.

Since LDTs are resistant to most classes of β-lactam antibiotics [63], crosslinking the cell wall via LDTs may contribute to the high antibiotic resistance to β-lactam antibiotics in the Rhizobiales. Here, we show that meropenem and faropenem inhibit ld- and dd-transpeptidation in *A. tumefaciens*, indicating that they may target LDTs and/or PBPs. Perhaps the loss of LDT activity disrupts the activity of high molecular weight PBPs if they function together in a growth pole complex. A combination of LDT and PBP-targeting antibiotics acted synergistically in killing *M. tuberculosis* [52], and a similar approach may also be effective against species in the Rhizobiales. Since faropenem causes swelling of the growth pole in *A. tumefaciens, S. meliloti* and *B. abortus* it is likely that the target(s) of this drug are conserved components of the growth machinery. Identification of the specific cellular target(s) of faropenem will provide candidate proteins for further characterization.

Expanding our understanding of the mechanism of polar growth in the Rhizobiales will help to shed light on the different strategies that can be employed by bacteria during cell elongation. Indeed, our findings highlight intriguing parallels with other, distantly related, polarly growing bacteria including the Actinobacteria. For example, PBP1a is essential in *Mycobacterium smegmatis* [64], an Actinobacteria that grows by bipolar elongation. In addition, PBP1a localizes to growth poles and is important for maintenance of rod shape in other Actinobacteria [65, 66]. Therefore, the reliance on PBP1a for synthesis of PG at the pole may be a key feature of polar-growing bacteria that arose independently in the Rhizobiales and Actinobacteria, indicating convergent evolution. Notably, the cell wall of polar-growing bacteria in both clades also contain a high proportion (~30-80%) of LDT-crosslinked PG [1, 67, 68], suggesting that ld-crosslinks may provide structural integrity. Furthermore, in *Mycobacterium* deletion of LDT-encoding genes leads to a loss of rod-shape [63], and LDTs also contribute to active PG synthesis of the sidewalls [37]. Finally, carbapenem antibiotics are routinely used to treat *M. tuberculosis* infections [20] hinting that the target of these drugs may be important in polar growing bacteria. Future work directed at characterizing the role of LDTs during polar growth in Rhizobiales and Actinobacteria is needed to determine if the high degree of ld-crosslinking is an innovation which allows for polar elongation to be adopted as the primary mode of growth. Overall, the possibility that there may be governing principles which allow for polar growth to emerge as a successful growth strategy is a fascinating concept which merits further study.

## Acknowledgements

We thank Zachary Taylor, Edward Hall, and Srinivas Tekkam for synthesis of cell wall probes and Gyanu Lamichhane for the generous gift of the penem and carbapenem antibiotics. We thank members of Jerod Skyberg’s lab for providing lab space and materials to work on *B. abortus*. We also thank members of the Brown lab and Marie Elliot for helpful discussions and critical reading of this manuscript. PB and MW were supported by the National Science Foundation, IOS1557806. MW was supported by the Life Sciences Fellowship at the University of Missouri. Research in the Cava lab (FC, AA) is supported by the Laboratory for Molecular Infection Medicine Sweden, the Knut and Alice Wallenberg Foundation, the Kempe Foundation and the Swedish Research Council. AA is supported by a MIMS/VR PhD position. EK was supported by the Life Sciences Research Foundation (LSRF) at the Harvard Medical School. AB received funding from the German Research Foundation (Project 269423233 - TRR 174). NIH grants R35GM122556 to YVB and GM113172 and to MSV and YVB and R35GM136365 to MSV. YVB is also supported by a Canada 150 Research Chair in Bacterial Cell Biology funded by the Canadian Institutes of Health Research.

## MATERIALS AND METHODS

### Bacterial strains, plasmids, and growth conditions

A list of all bacterial strains used in this study is provided in Supplemental Table 1. *Agrobacterium tumefaciens* C58 and derived strains were grown in ATGN minimal media [69] without exogenous iron at 28°C with shaking. When appropriate, kanamycin (KAN) was used at the working concentration of 300 μg/ml. When indicated, isopropyl β-D-1-thio-galactopyranoside (IPTG) was used as an inducer at a concentration of 1 mM. *Sinorhizobium meliloti* strains were grown in Tryptone-Yeast (TY) medium. When appropriate, KAN was used at the working concentration of 100 μg/ml, Gentamicin (GM) was used as 20 μg/ml, and IPTG was used at a concentration of 500 μg/ml. *Brucella abortus* strain S19 was grown in brucella broth. *E. coli* strains were grown in Luria-Bertani medium at 37°C. For *E. coli* DH5α and S17-1 λ *pir*, when appropriate 50 μg/ml or 30 μg/ml of KM, respectively, was added.

### Construction of strains and plasmids

A list of all primers used in this study is provided in Supplemental Table 2. For amplification of target genes, primer names indicate the primer orientation and added restriction sites. All expression vectors were verified by sequencing. All vectors were introduced into *A. tumefaciens* strains utilizing standard electroporation protocols [70] with the addition of IPTG in the media when introducing plasmids into depletion backgrounds.

### Construction of deletion/depletion plasmids and strains

Vectors for gene deletion by allelic exchange were constructed using recommended methods for *A. tumefaciens* [70]. Briefly, 500-bp fragments upstream and 500 bp downstream of the target gene were amplified using primer pairs P1/P2 and P3/4 respectively. Amplicons were spliced together by SOEing using primer pair P1/P4. The amplicon was digested and ligated into pNTPS139. The deletion plasmids were introduced into *A. tumefaciens* by mating using an *E. coli* S17 conjugation strain to create KM resistant, sucrose sensitive primary integrants. Primary integrants were grown overnight in media with no selection. Secondary recombinants were screened by patching for sucrose resistance and KM sensitivity. Colony PCR with primers P5/P6 for the respective gene target was used to confirm deletion. PCR products from P5/P6 primer sets were sequenced to further confirm deletions.

For depletion strain construction, target genes (*pbp1a* or *pbp3a*) were amplified, digested and ligated into pUC18-mini-Tn*7*T-GM-P_lac_. The mini-Tn*7* vector, along with the pTNS3 helper plasmid, were introduced into C58Δ*tetRA::a-att*Tn7 as described previously [27]. Transformants were selected for GM resistance and insertion of the target gene into the a-*att* site was verified by colony PCR using the tet forward and Tn7R109 primer. PCR products were sequenced to confirm insertion of the correct gene. Next, the target gene was deleted from the native locus as described above in the presence of 1 mM IPTG to drive expression of the target gene from the engineered site.

To generate the *S. meliloti* strain lacking the five non-essential PBPs, the corresponding genes were consecutively deleted from the Rm2011 *rgsP-egfp* genome using the sucrose selection method. [71] To generate the *S. meliloti* MrcA1 depletion strain, first plasmid pK18mobsac-mrcA1del was integrated into the Rm2011 *rgsP-egfp* genome, then an ectopic *mrcA1* copy was introduced on plasmid pGCH14-mrcA1 followed by sucrose selection of mutant clones with deletion of the native *mrcA1* allele. The curable plasmid pGCH14, which is maintained in single copy in *S. meliloti* due to the replication operon *repABCpMlb_lacO*, prone to repression by LacI was used as vector to conditionally establish the ectopic copy of *mrcA1* under control of its native promoter. pSRKKm, carrying *lacI*, was introduced into the strain with chromosomal deletion of *mrcA1*, carrying pGCH14-mrcA1, and the resulting strain Rm2011 *rgsP-egfp mrcA1*^dpl^ was grown in presence of 500 μm IPTG. Growth in the absence of IPTG induced MrcA1 depletion due to loss of the ectopic *mrcA1* copy.

### Phase and Fluorescence microscopy

A small volume (~1 μl) of cells in exponential phase (OD_600_ = 0.2 - 0.4) was applied to a 1% ATGN agarose pad as described previously [72]. DIC, Phase contrast and epifluorescence microscopy were performed with an inverted Nikon Eclipse TiE and a QImaging Rolera em-c2 123 1K EMCCD camera with Nikon Elements Imaging Software. For time-lapse microscopy, images were collected every ten minutes, unless otherwise stated.

### Quantification of cell length distributions

Cells were grown overnight in ATGN. Cells were diluted in ATGN to an OD_600_ = 0.2 and allowed to grow until reaching an OD_600_ = 0.4 - 0.6. Live cells were imaged using phase contrast microscopy and cell length distributions of the indicated number of cells per strain were determined using the longest medial axis as measured by MicrobeJ software [24].

### Quantification of cell morphologies, FDAA and FDAAD labeling patterns

For *A. tumefaciens*, cells were grown overnight in ATGN media and diluted in the same conditions to an OD_600_ = 0.20 and allowed to grow until reaching an OD_600_ = 0.4 - 0.6. At this point cells were labeled with 1mM of the fluorescent-d-amino acid (FDAA) HCC-amino-d-alanine (HADA) or the fluorescent dipeptide (FDAAD) NBd-amino-d-alanine-d-alanine (HADA—DA) as previously described [36, 38, 41]. Immediately following a five-minute incubation, cells were ethanol fixed to prevent further growth. Phase contrast and epifluorescence microscopy was performed on the reported number of cells. For *E. coli*, cells were grown overnight at 37°C in M9 + 0.2% glucose minimal medium and diluted in the same conditions to an OD_600_ = 0.1 and allowed to grow until reaching an OD_600_ = 0.4 - 0.6. At this point cells were labeled with 1 mM of HADA—DA. After an incubation of 90 minutes cells were ethanol fixed and washed 3 times in 1 mL PBS before imaging. For *B. subtilis 3610*, cells were grown overnight at 37°C in S750 + 1% glucose defined minimal medium and diluted in the same conditions to an OD_600_ = 0.1 and allowed to grow until reaching an OD_600_ = 0.4 - 0.6. At this point cells were labeled with 5 mM of HADA—DA. After an incubation of 120 minutes cells were ethanol fixed and washed 3 times in 1 mL PBS before imaging. For *S. venezuelae*, cells were grown overnight at 30°C in LB medium and diluted in the same conditions to an OD_600_ = 0.1 and allowed to grow until reaching an OD_600_ = 0.4 - 0.6. At this point cells were labeled with 1 mM BODIPY FL-amino-d-alaninyl-D-alanine (BADA—DA). After an incubation of 15 minutes, cells were washed once with 1 mL LB and resuspended in 500 uL LB containing 2 mM HADA—DA. After an incubation of 15 minutes, cells were washed once with 1 mL LB and resuspended in 500 uL LB containing 1 mM ATTO 610-amino-d-alaninyl-D-alanine (Atto610ADA—DA). After an incubation of 15 minutes cells were ethanol fixed and washed 3 times in 1 mL PBS before imaging.

Demographs were constructed using MicrobeJ. For demographs, cells were arranged from top to bottom according to their cell lengths and each cell was oriented such that the new pole (defined as the cell pole with the higher fluorescence intensity as determined by FDAA or FDAAD labeling or the smaller pole diameter in cells without label) was oriented to the right.

### Synthesis of HADA—DA

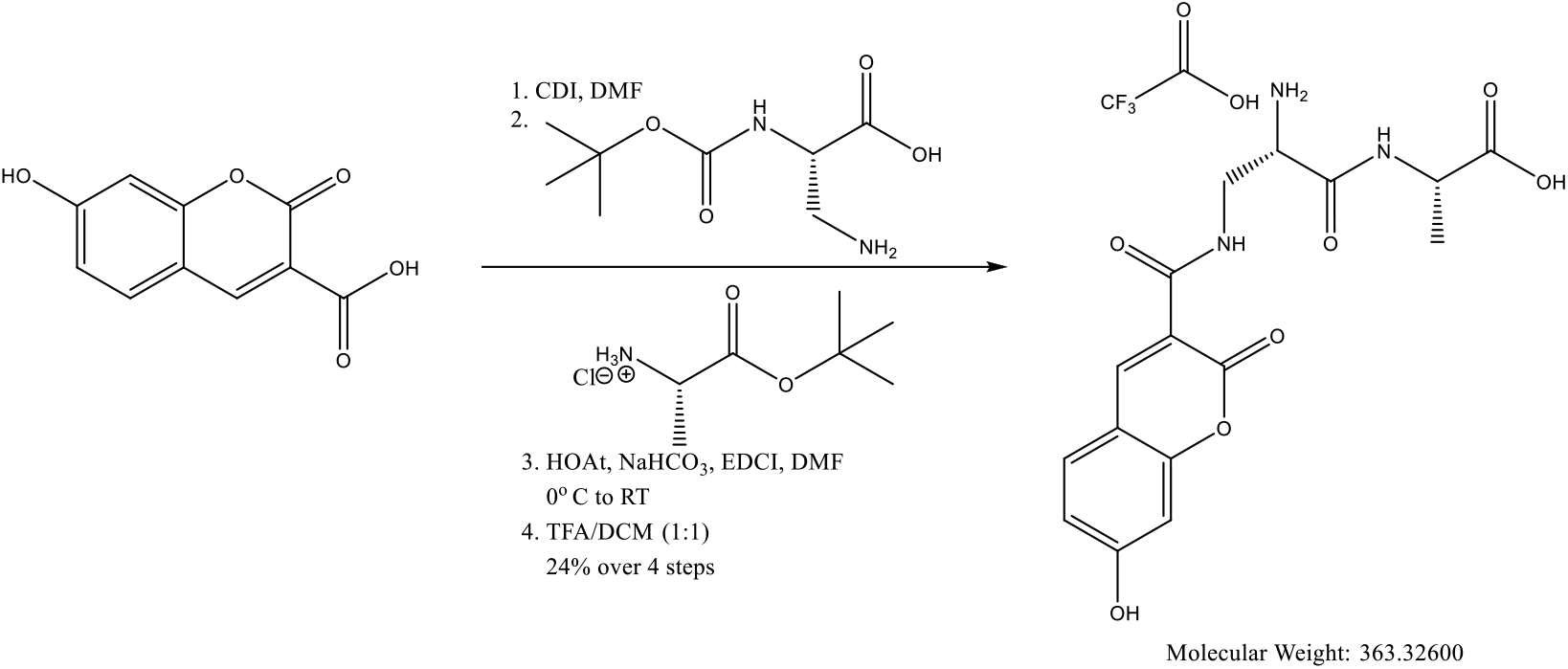

A solution of 7-hydroxycoumarin-3-carboxylic acid (HCC) (105 mg, 0.51 mmol) and carbonyldiimidazole (83 mg, 0.51 mmol) in anhydrous DMF (5 mL) was stirred at room temperature (RT) for 2 h under an atmosphere of argon. Boc-d-2,3-diaminopropionic acid (104 mg, 0.51 mmol) was then added and the reaction mixture was allowed to stir at RT for 18 h. DMF was removed *in vacuo*. The residue was diluted with EtOAc (35 mL), washed with 1N HCl (20 mL) and water (30 mL). The combined aqueous layers are back-extracted with EtOAc (20 mL). The combined organic layers were washed with brine (20 mL), dried over sodium sulfate, filtered, and concentrated to dryness. The crude product was dissolved in anhydrous DMF. d-alanine tert-butyl ester hydrochloride (111 mg, 0.612 mmol), HOAt (83.5 mg, 0.612 mmol) and NaHCO_3_ (94 mg, 1.12 mmol) were added successively, and the reaction mixture was cooled to 0 °C. EDCI (117 mg, 0.612 mmol) was then added, and the reaction mixture was stirred for 12 h. The solvent was removed in *vacuo* and the product was diluted with EtOAc, washed with 1N HCl, water, brine, dried over sodium sulfate, filtered and concentrated. The crude product was dissolved in DCM/TFA (1:1, 4 mL) mixture, stirred for 1 h and evaporated to dryness. The yellow color solid was dissolved in acetonitrile/water and purified by reverse phase HPLC and lyophilized to yield the desired product as a pale yellow solid (63 mg, 26%). 1H NMR (500 MHz, DMSO-*d6*) 11.37 (br s, 1H), 8.91 (m, 1H), 8.90 (m, 1H), 8.81 (s, 1H), 8.42 (br s, 3H), 7.83 (m, 1H), 5.94 (m, 1H), 5.89 (s, 1H), 4.24 (m, 1H), 4.04 (br s, 1H), 3.77 (m, 1H), 3.68 (m, 1H), 1.32 (s, 1H); HRMS-ESI-TOF m/z calc C16H18N3O7 (M+H) 364.1145, Found 364.1161.

### PG compositional analysis

For PG analysis, three cultures of each of strain were grown overnight in 3 ml culture tubes of ATGN minimal media at 28°C with shaking; the + PBP1a strain was supplemented with 1mM IPTG. The 3 ml cultures were then added to 50 ml flasks of fresh ATGN and allowed to grow under the same conditions until reaching an exponential phase OD_600_ of 0.5-0.6. Cells were then pelleted by centrifugation at 4000 x *g* for 10 minutes. Cell pellets were washed three times with ATGN by centrifugation and resuspension to remove IPTG. After the final wash, the 3 cell pellets from the + PBP1a strain were split and resuspended in 50 ml ATGN with or without IPTG. Each culture was grown for 16 hours (hr). to an OD_600_ of 0.6 After 146 h of growth, 50 ml of the exponential cultures were collected and pelleted by centrifugation at 4000 x *g* for 20 minutes. Cell pellets were resuspended in 3 mL of ATGN and 6 mL of 6% SDS and stirred with magnets while boiling for 4 h. Next, samples were removed from heat but continued to stir overnight. Samples were then shipped to Cava laboratory for purification and analysis. Upon arrival, cells were boiled and simultaneously stirred by magnets for 2 h. After 2 h, boiling was stopped, and samples were stirred overnight. PG was pelleted by centrifugation for 13 minutes (min) at 60,000 rpm (TLA100.3 Beckman rotor, Optima Max-TL ultracentrifuge; Beckman), and the pellets were washed 3 to 4 times by repeated cycles of centrifugation and resuspension in water. The pellet from the final wash was resuspended in 50 μl of 50 mM sodium phosphate buffer, pH 4.9, and digested overnight with 100 μg/ml of muramidase at 37°C. Muramidase digestion was stopped by boiling for 4 min. Coagulated protein was removed by centrifugation for 15 min at 15,000 rpm in a desktop microcentrifuge. The muropeptides were mixed with 15 μl 0.5 M sodium borate and subjected to reduction of muramic acid residues into muramitol by sodium borohydride (10 mg/ml final concentration, 20 min at room temperature) treatment. Samples were adjusted to pH 3 to 4 with orthophosphoric acid and filtered (0.2-μm filters). Analysis of muropeptides was performed on an ACQUITY Ultra Performance Liquid Chromatography (UPLC) BEH C18 column, 130Å, 1.7 μm, 2.1 mm x 150 mm (Water, USA) and detected at Abs. 204 nm with ACQUITY UPLC UV-visible detector. For data shown in Figure 4B and Supplementary Figure 7B, muropeptides were separated with organic buffers at 45°C using a linear gradient from buffer A (formic acid 0.1% (v/v) in water) to buffer B (formic acid 0.1% (v/v) in acetonitrile) in a 18 minutes run with a 0.25 ml/min flow. For data shown in Figure 5A and Supplementary Figure 5A, muropeptides were separated using a linear gradient from buffer A (sodium phosphate buffer 50 mM pH 4.35) to buffer B (sodium phosphate buffer 50 mM pH 4.95 methanol 15% (v/v)) with a flow of 0.25 mL/min in a 20 min run. Individual muropeptides were quantified from their integrated areas using samples of known concentration as standards. Muropeptide abundance was statistically compared using a one-way ANOVA with Tukey’s multiple comparisons test.

## Notes

### Competing Interest Statement

The authors have declared no competing interest.

