## Supplementary Materials for "Unipolar peptidoglycan synthesis in the Rhizobiales requires an essential class A penicillin-binding protein"

**1) SUPPLEMENTARY FIGURES, LEGENDS, and MOVIE LEGENDS**
**2) SUPPLEMENTARY METHODS**
**3) SUPPLEMENTARY TABLES S1 and S2**

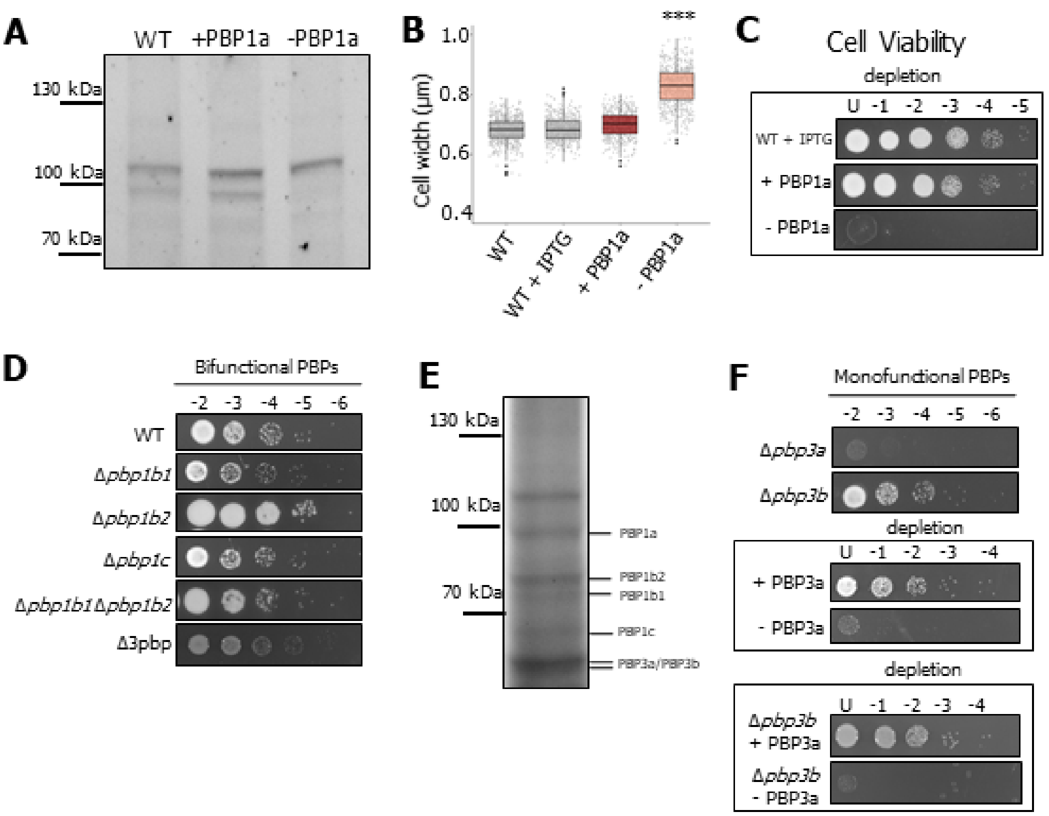

**Supplementary Figure 1. Characterization of PBP mutants.** (**A**) Bocillin labeling of cell membrane preps of wildtype (WT) and the PBP1a depletion strain. The PBP1a depletion strain was grown in the presence (+ PBP1a) or absence of 1mM IPTG for 16 hours (- PBP1a). The band corresponding to PBP1a protein is not detected after 16 hours of depletion. (**B**) Cell width distributions of wildtype cells compared to the PBP1a depletion strain grown with or without IPTG for 16 hours. The indicated strains were grown as in Figure 1 and subjected to cell width measurements using MicrobeJ [24]. The data are shown as box and whisker plots in the style of Tukey [25]. Distributions of cells significantly different from WT are indicated (***; One-Way ANOVA with Bonferroni correction, p< 2*10^16). n = > 800 per strain. (**C**) Spot assay for cell viability of the PBP1a depletion strain. Images are representative of three independent biological replicates. Dilutions are indicated above each spot and depletion strains are indicated in the boxes. (**D**) Spot assays for cell viability of bifunctional PBP deletion strains. Dilutions are indicated above each spot, and images are representative of three independent biological replicates. (**E**) Bocillin labelling of wildtype cell membrane prep., which is a representative image of two independent experiments. Each PBP is labeled according to its predicted molecular weight. (**F**) Spot assays for cell viability of monofunctional PBP deletion and depletion strains. Dilutions are indicated above each spot and depletion strains are indicated in the boxes. Images are representative of three independent biological replicates.

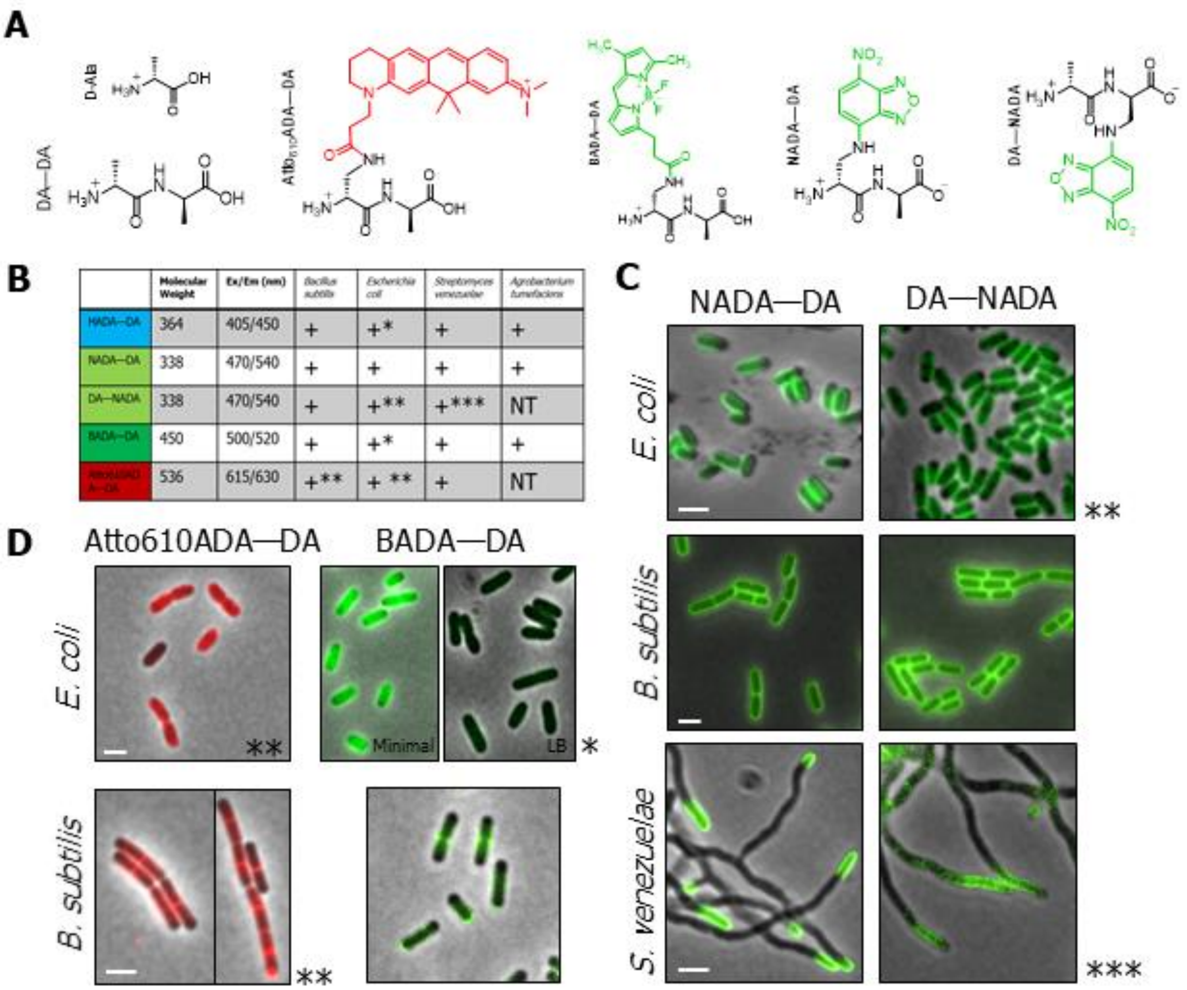

**Supplementary Figure 2. Qualitative evaluation of FDAAD labeling in different bacteria.** (**A**) Structures of the **f**luorescent **D** -**a**mino **a**cid **d**ipeptides (FDAADs) investigated in this work. (**B**) Table indicating the labelling pattern of different fluorescent D-amino acid dipeptides (FDAADs) in different bacterial species. BADA—DA is a brighter fluorescent alternative to NADA—DA in most cases. +**:** Significant labeling (signal-to-background >1) NT: Not tested * Indicates where a quantitative comparison between minimal media, e.g. M9, and rich media, e.g. LB, where minimal media showed stronger labeling, ** patchy or peculiar labeling patterns different from FDAA labeling. *** Indicates where a quantitative comparison between DA—NADA and NADA—DA is done and labeling shows weaker DA—NADA labeling, likely due to the loss of the fifth position NADA upon crosslinking by PBPs. (**C**) Merged phase and fluorescent images of representative cells comparing labeling between NADA—DA and DA-NADA in the species listed. (**D**) Merged phase and fluorescent images of representative cells comparing labeling between Atto610ADA—DA and BADA—DA. All scale bars are 2 µm.

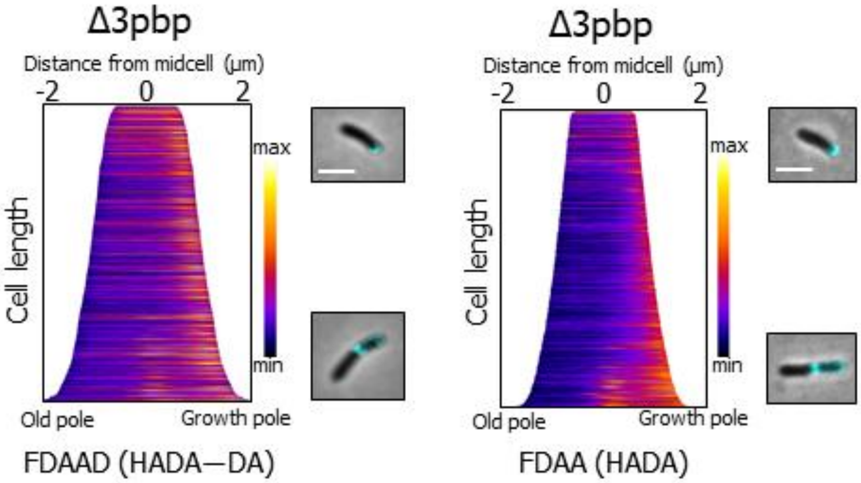

**Supplementary Figure 3. Comparison of the labelling patterns of different fluorescent cell wall probes in the Δpbp3 strain.** (**A**) Demographs depict incorporation of either FDAADs or FDAAs at a population level. Median profiles of the fluorescence channel of more than 1200 cells per strain are stacked and ordered by cell length. Merged phase and fluorescent channels of cells with representative polar and septal labeling of fluorescent probes are shown beside each demograph. Scale bar: 2 µm.

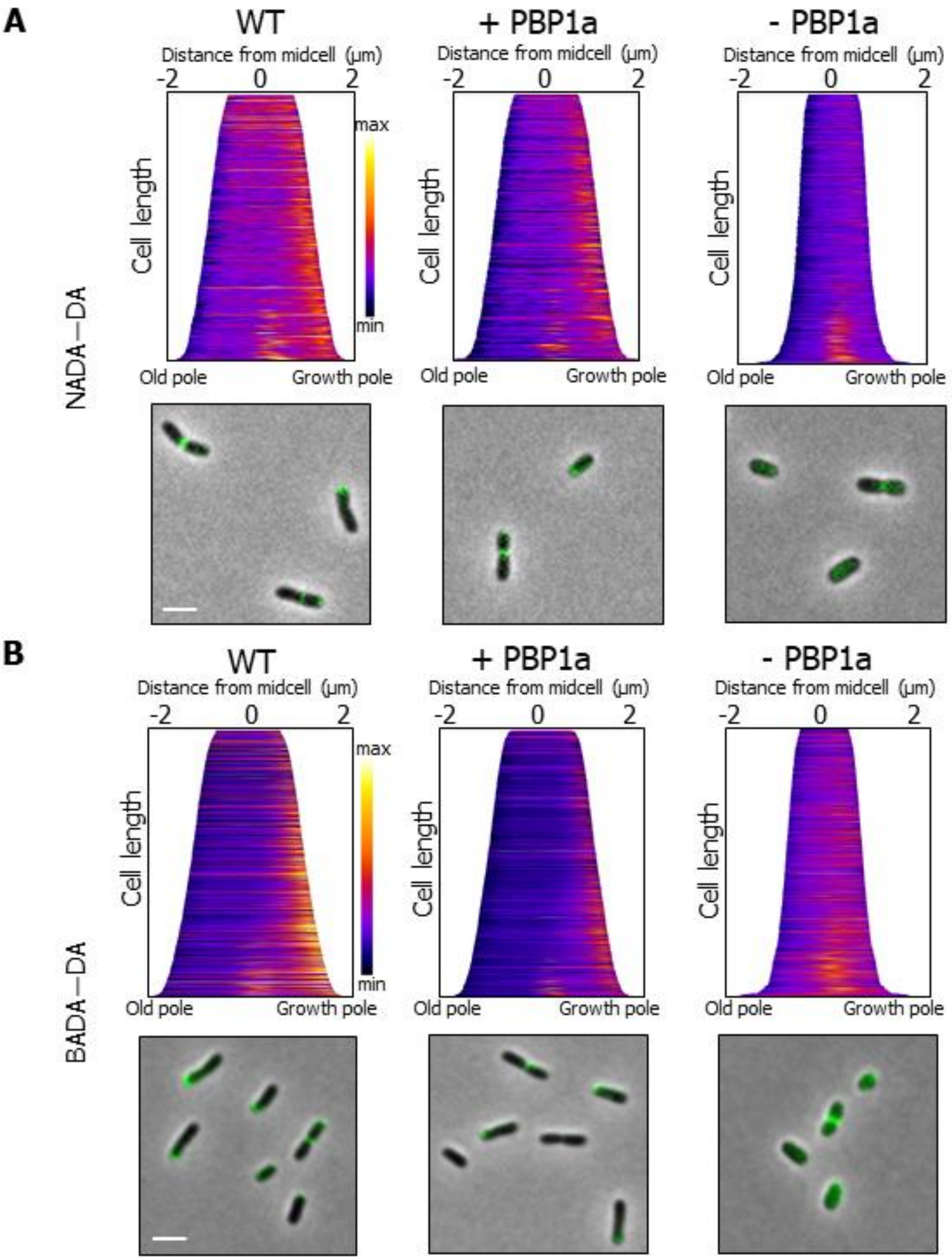

**Supplementary Figure 4. Comparison of the labelling patterns of NADA—DA and BADA—DA.** (**A**) Demographs depict incorporation of NADA—DA at a population level. Median profiles of the fluorescence channel of more than 400 cells per strain are stacked and ordered by cell length. Merged phase and fluorescent channels of cells with representative polar and septal labeling of dipeptides are shown below each demograph. (**B**) Demographs depict incorporation of BADA—DA at a population level. Median profiles of the fluorescence channel of more than 400 cells per strain are stacked and ordered by cell length. Merged phase and fluorescent channels of cells with representative polar and septal labeling of dipeptides are shown below each demograph. Scale bar: 2 µm.

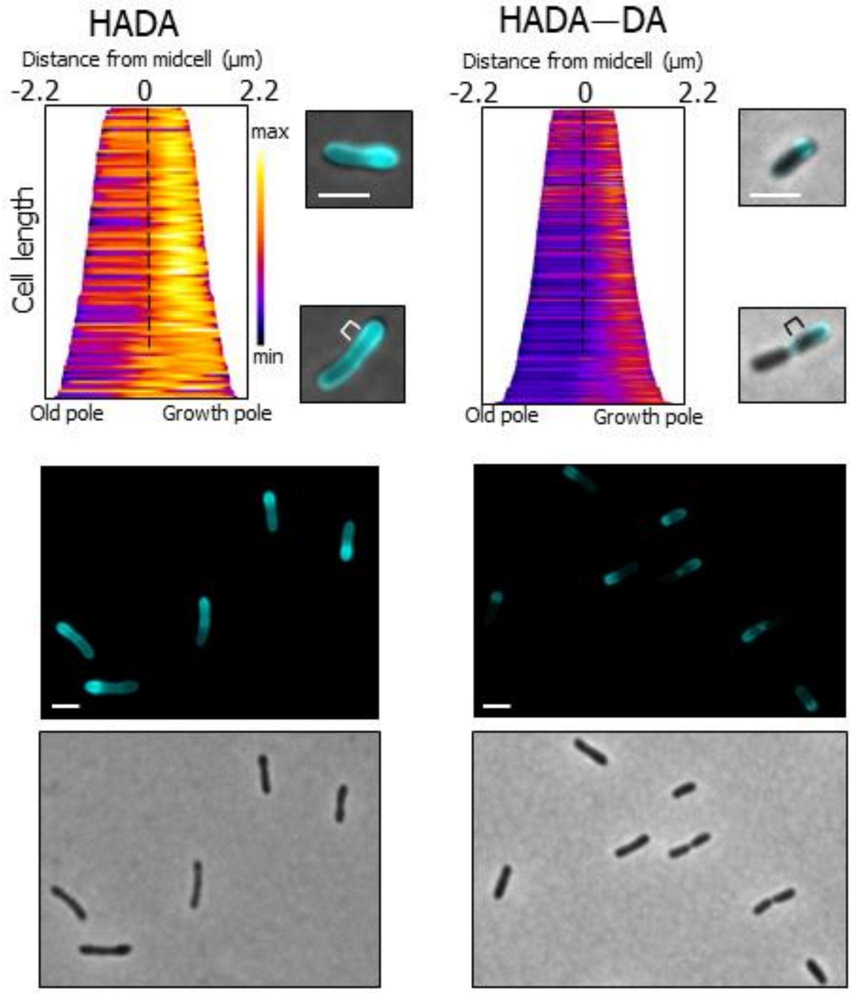

**Supplementary Figure 5. Comparison of the labelling pattern after a long pulse with FDAAs and FDAADs.**(**A**) Demographs depict incorporation of either HADA or HADA—DA at a population level. Median profiles of the fluorescence channel of 95 cells for HADA and 190 cells for HADA—DA are stacked and ordered by cell length. Scale bar for the HADA demograph represents intensity and ranges from 900-4800 a.u. Scale bar for the HADA—DA demograph ranges from 100-900 a.u. Merged phase and fluorescent channels of cells with representative polar and septal labeling of dipeptides are shown beside each demograph. Brackets point to the sidewalls of the new pole. Below the demographs are images of several representative cells, the fluorescent and phase are shown separately Scale bar: 2 µm.

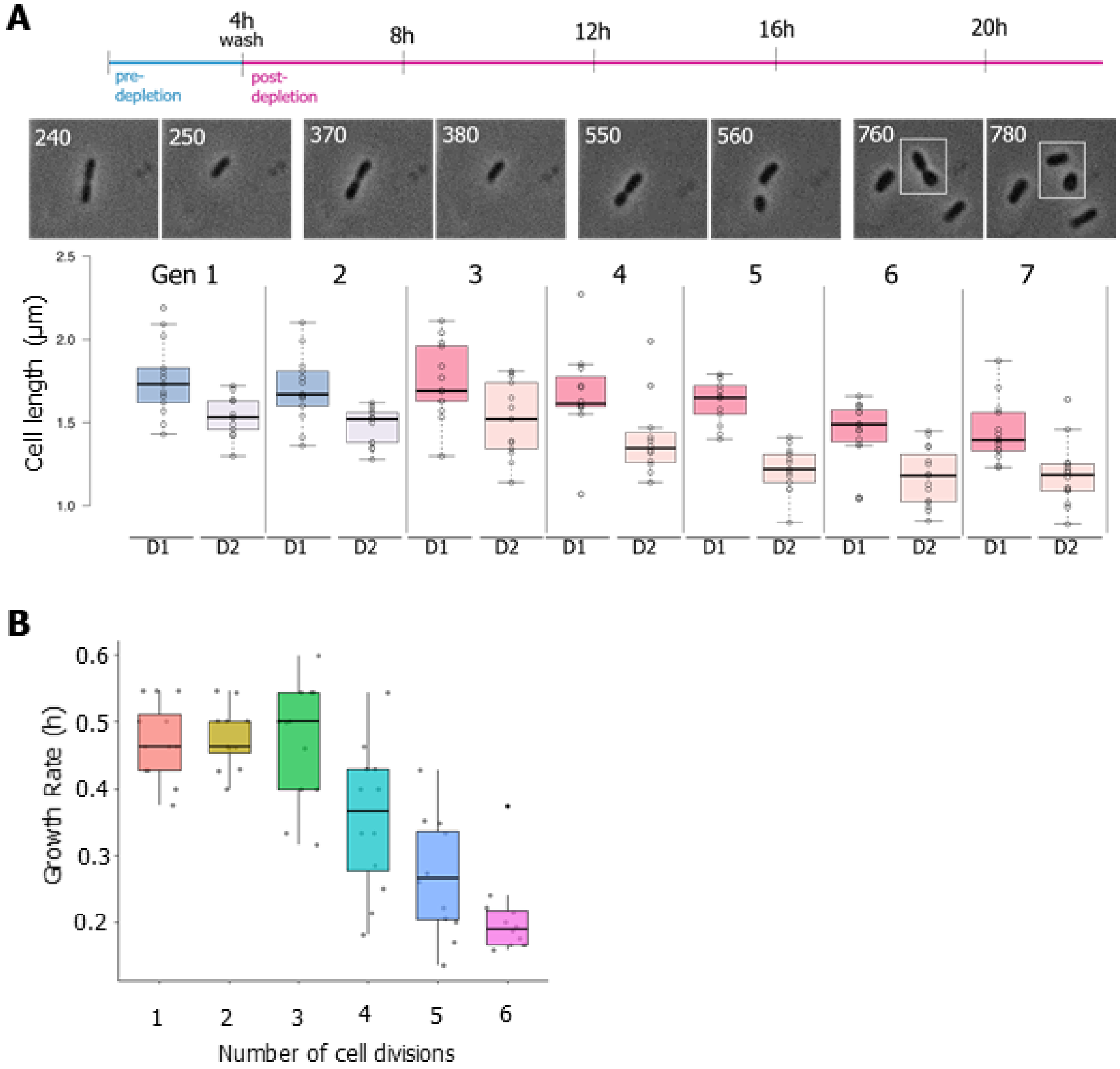

**Supplementary Figure 6. Impacts of the PBP1a depletion accumulate over multiple generations.** (**A**) Shown are cell length distributions of the PBP1a depletion strain grown in a microfluidic device. A timeline of the experiment is provided. Cells were induced with 1mM IPTG for 4 hours before washing to deplete the cells of PBP1a for the remainder of the experiment. Twelve cells were followed for 7 generations, corresponding to 24 hours of growth in the device and the cell length of the two daughter cells was measured one frame after the cell divided (D1 and D2 for daughter cell 1 and daughter cell 2). D1 was always designated as the cell that remained attached to the microfluidic device, or the old pole daughter cell. Rarely, if D2 (the new pole daughter cell) washed away before it could be measured, the cell length was calculated by subtracting the length of D1 from the mother cell length just before it divided. Cell lengths measurements were collected using MicrobeJ [24]. The data are represented as box and whisker plots in the style of Tukey [25]. Phase images of one representative cell just before cell division and just after cell division from four different generations (gen) corresponding to pre-depletion, early, mid and late depletion timepoints. Time is indicated in minutes. (**B**) Growth rate in hours (h) of the twelve cells in A was calculated by taking the inverse of the time between cell divisions.

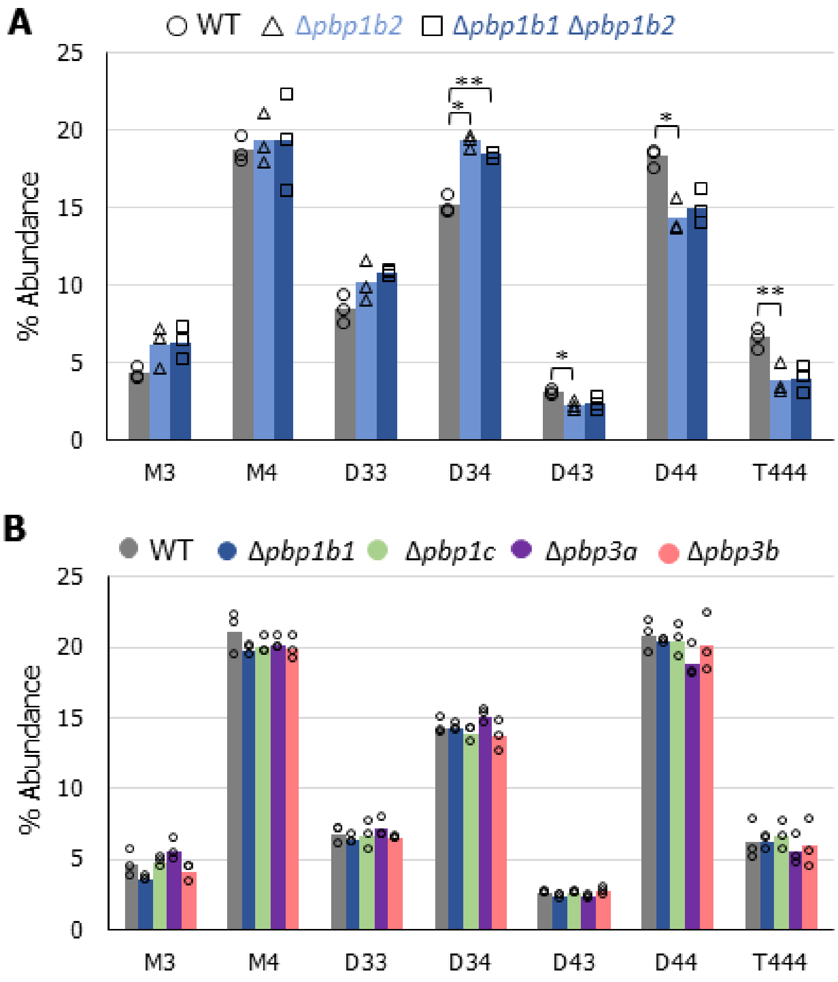

**Supplementary Figure 7. Peptidoglycan composition of PBP mutants.** (**A**) Bar graphs showing the average abundance of muropeptides obtained by UPLC analysis from the indicated strains. Data shown are averages taken from analysis of three independent biological samples. Samples significantly different are indicated (One-Way ANOVA with Tukey's multiple comparison test, * $p<0.05$, ** $p<0.005$). (**B**) Bar graphs showing the average abundance of muropeptides obtained by UPLC analysis from the indicated strains. Data shown are averages taken from analysis of three independent biological samples. The difference between samples was not significant (One-Way ANOVA with Tukey's multiple comparison test).

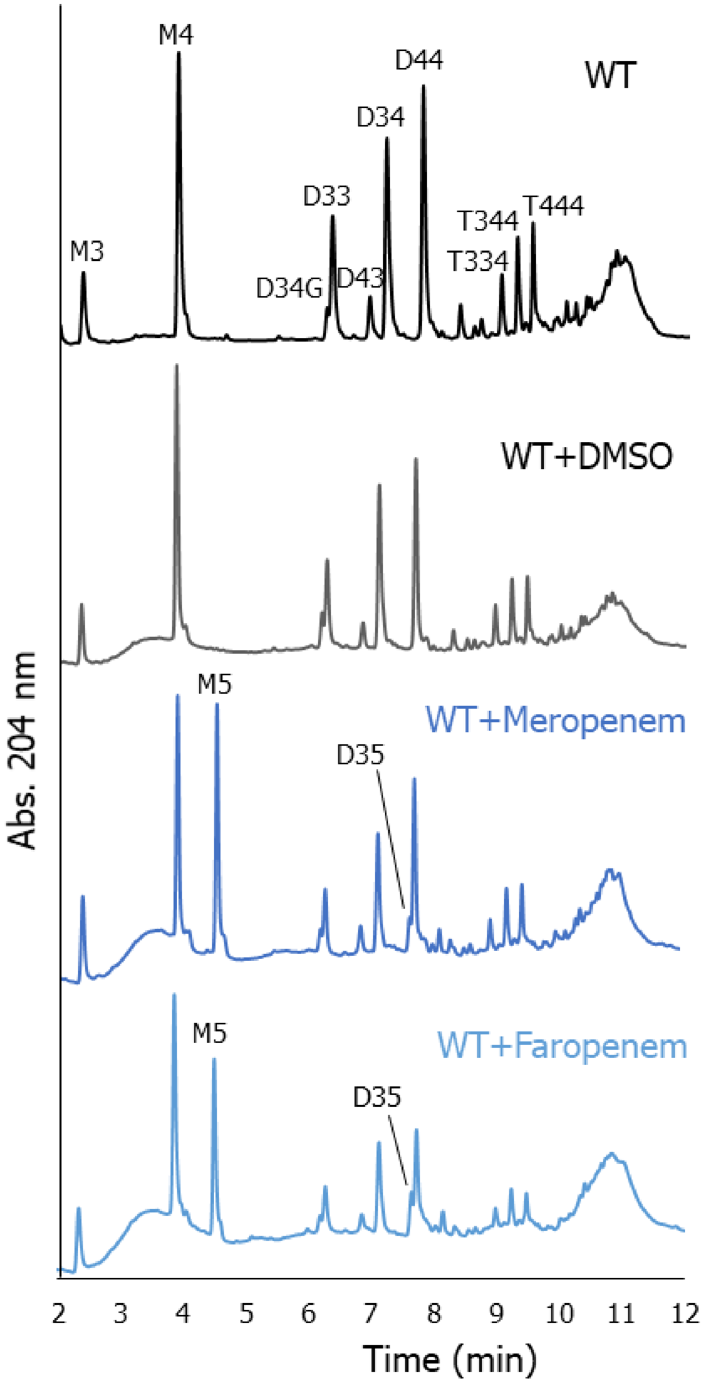

**Supplementary Figure 8. Impact of antibiotic treatment on peptidoglycan composition of *A. tumefaciens.*** (**A**) Representative UPLC profiles showing the abundance of muropeptides obtained from the indicated strains. Cells were grown in the presence of DMSO or 1.5 µg/mL faropenem for 6 hours, corresponding to the onset of polar swelling induced by faropenem treatment before harvesting PG. Cell were grown in the presence of 1.5 µg/mL meropenem for 4 hours, corresponding to the onset of mid-cell swelling induced by meropenem treatment before harvesting PG.

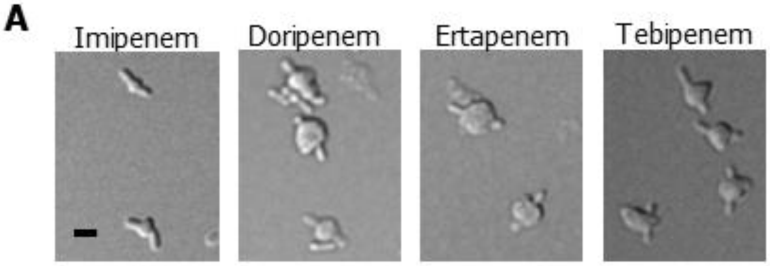

**Supplementary Figure 9. Morphology of carbapenem treated cells.** (**A**) Representative images of wildtype cells grown in the presence of 1.5 µg/mL imipenem, doripenem, and ertapenem and 50 µg/mL tebipenem. Cells were incubated with antibiotics for 24 hours, then spotted on a 1% agarose pad and imaged using DIC microscopy. Scale bar: 2 µm

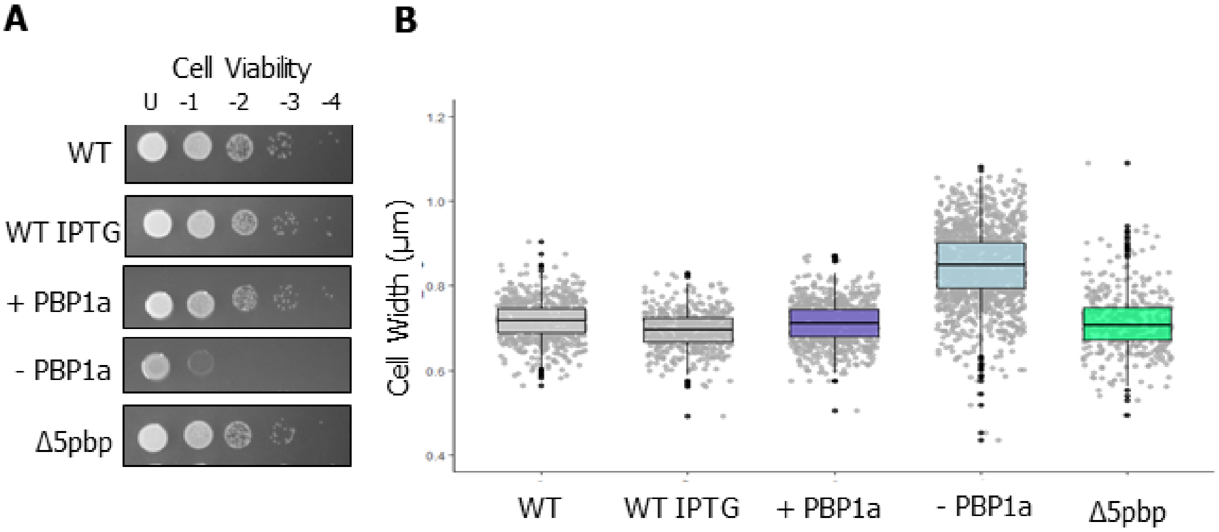

**Supplementary Figure 10. Characterization of *S. meliloti* PBP strains.** (**A**) Spot assays for cell viability of PBP deletion and depletion strains of *S. meliloti*. Dilutions are indicated above each spot, and images are representative of three independent biological replicates. (**B**) Cell width distributions of all strains from figure 6B. The indicated strains were subjected to cell width measurements using MicrobeJ [24]. The data are shown as box and whisker plots in the style of Tukey [25]. n = >400 per strain.

**Supplementary Movie 1. Cells lacking both PBP3a and PBP3b are blocked for cell division.** DIC image series of the PBP3b deletion PBP3a depletion strain. Cells were pre-depleted of IPTG for 4 hours before spotting onto a 1% ATGN agar pad. Images were acquired every 10 minutes, and the movie is played at 10 frames per second for a total of 130 frames

**Supplementary Movie 2. PBP1a depleted cells get successively smaller and rounder over multiple generations in a microfluidic device.** Phase image series of *the* PBP1a depletion strain growing in ATGN minimal medium in a microfluidic device. Cell were grown in the presence of IPTG to induced PBP1a expression for 4 hours, washed of IPTG and grown without inducer for the remainder of the video. Images were acquired every ten minutes, and the movie is played at 10 frames per second for a total of 130 frames.

**Supplementary Movie 3. Faropenem causes swelling of the growth pole.** DIC image series of wildtype *A. tumefaciens* cells growing on a 1% ATGN agar pad supplemented with 1.5 µg/mL faropenem. Images were acquired every 10 minutes, and the movie is played at 10 frames per second for a total of 130 frames.

### SUPPLEMENTARY METHODS

**Spotting Assays** For cell viability spot assays, exponentially growing cultures were diluted to $OD_{600}$ = 0.05 and serially diluted in ATGN. (this is the undiluted cells – U). 4 µl of each dilution was spotted onto an ATGN plate with or without IPTG and incubated at 28°C for 2 days before imaging.

**Bocillin-FL labeling of Membrane Extract** Cells were grown from a single colony in a 3 mL of ATGN media overnight, then the 3 mL was transferred to a 50 mL flask grown overnight, and sub-cultured in 1L of ATGN media the next morning and grown to an $OD_{600}$ of 1.0. To extract the cell membrane, cells were pelleted at 4,400 x g for 10 minutes at 4 °C, resuspended in buffer (10 mM potassium phosphate + 140 mM NaCl, pH 7.0), and then sonicated. After sonication cells were passed through an 20G needle 10 times to further lyse the cells. Centrifugation cell lysate at 12,000 x g for 10 min at 4 °C. Centrifuge the supernatant fractions at 150,000 x g for 40 min at 38,000 rpm. Resuspend pellets in 1 mL buffer (10 mM potassium phosphate + 140 mM NaCl, pH 7.0). Total protein concentration was measured by Bradford assay and 0.7 µg/mL of the preparation was incubated in the presence of buffer (20 mM potassium phosphate + 140 mM NaCl) and 0.1 mM Bocillin-FL (Thermo Fisher Scientific, Waltham, MA) for 5 minutes at 37 °C in the dark, and then the reaction was quenched upon addition of Laemmli buffer to 1x final concentration. Samples were boiled for 5 minutes, then 30 µL was separated on a 4-12% SDS-PAGE gradient gel. Gels were imaged on BioRad gel image (Alexa488 setting).

**Microfluidics** The CellASIC™ ONIX Microfluidic Platform was used with CellASIC ONIX plate for bacteria cells (4 chamber) Cat number: B04A-03-5PK. The device was set up according to the CellASIC® ONIX B04A-02 Microfluidic Bacteria Plates User Guide in conjunction with an inverted Nikon Eclipse TiE and a QImaging Rolera em-c2 123 1K EMCCD camera with Nikon Elements Imaging Software. Images were taken with high-numerical aperture (NA) objectives in the analysis channel at 60x magnification every 10 minutes. The microfluidic device was inoculated by pumping +PBP1a cells from an $OD_{600}$ 0.1 culture in the loading chamber according the user manual. ATGN media + IPTG was flowed over cells at a low pounds per square inch (psi) for 1 hour. During this time, permanent adhesion of the bacteria to the microchannel wall was observed. Device seeding was terminated after 1 h, and ATGN medium was pumped through the incubation region at a higher psi for 5 minutes to flush away any unadhered cells. The cells in the incubation region were grown for 4 hours with ATGN + IPTG media, then ATGN media without IPTG was flowed in at a higher psi for 5 minutes to quickly wash away the ITPG. The psi was returned to the normal rate for the remaining 20 hours.

**FDAAD Synthesis**

**BADA—DA:**

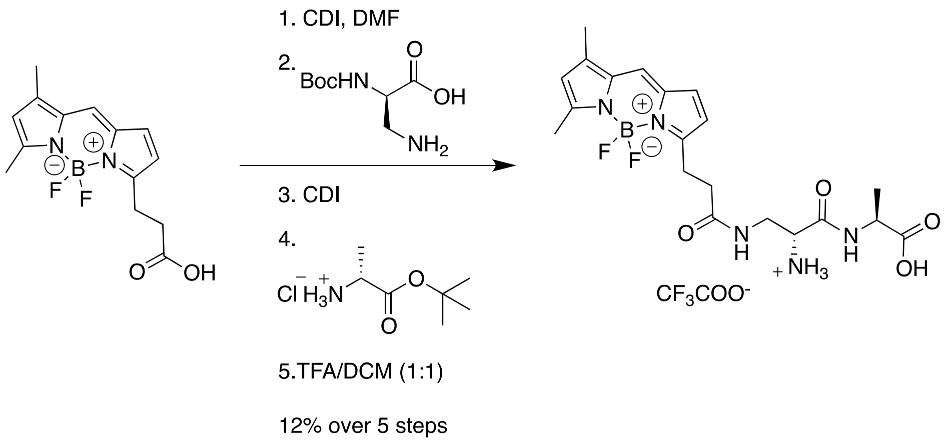

EverFluor FL acid (known as BODIPY FL® TM of Molecular probes) was purchased from Setareh Biotech, LLC. Carbonyldiimidazole (CDI) was purchased from Sigma Aldrich and used as received.

Under a blanket of argon, EverFluor FL acid (5.0 mg, 0.017 mmol) was dissolved in DMF (0.2 mL). To this stirring solution was added CDI (3.2 mg, 0.020 mmol) at room temperature. After 3 hours of stirring, Boc-D-2,3-diaminopropionic acid was added (3.6 mg, 0.018 mmol). The reaction was allowed to continue overnight. CDI (4.7 mg, 0.029 mmol) was then added to this reaction mixture. After 3 hours, D-AlaOtBu·HCl salt (5.0 mg, 0.028 mmol) was added. Following stirring for ~24 hours, the reaction was concentrated *in vacuo.* The crude product was then diluted in a TFA/DCM mixture at room temperature (1:1, ~5 mL). After deprotection (~ 5 hours) the solvent was removed *in vacuo* and the crude product was isolated by reverse phase HPLC (30-90% ACN/$H_2O$ over 10 minutes, RT = 5.7 minutes) and freeze dried to provide an orange powdery solid (0.9 mg, 0.002 mmol, 12%). $^1$H NMR (600 MHz, Acetonitrile-d3) δ 7.69 (s, 1H), 7.58 (d, J = 6.9 Hz, 1H), 7.37 (s, 1H), 7.00 (d, J =3.8 Hz, 1H), 6.32 (d, J = 3.9 Hz, 1H), 6.22 (s, 1H), 4.40 (t, J = 7.3 Hz, 1H), 4.13 (s, 1H), 3.82 – 3.68 (m, 1H), 3.62 (d, J =

14.4 Hz, 1H), 3.15 (t, J = 7.8 Hz, 2H), 2.60 (t, J = 7.2 Hz, 2H), 2.50 (s, 3H), 2.25 (s, 4H), 1.39 (dd, J = 7.2, 2.1 Hz, 3H). HRMS-ESI-TOF m/z calc 450.2128, Found 450.2106 $[M+H]^+$

**ATTO-610ADA—DA:**

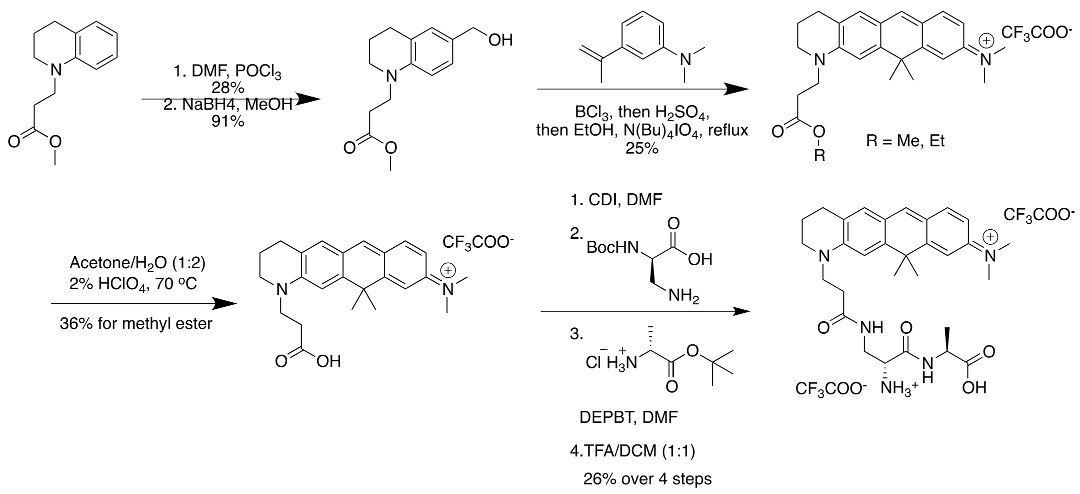

**Methyl 3-(6-formyl-3,4-dihydroquinolin-1(2*H*)-yl)propanoate:**

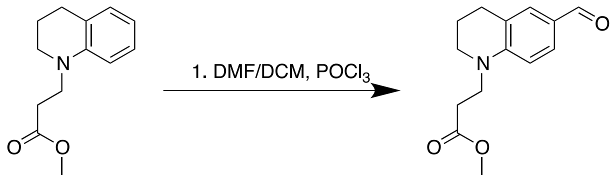

To a stirring solution of substrate (19.68 grams, 89.8 mmol) in a DCM/DMF mixture (28.0 mL DMF, circa 150 mL DCM) at 0 $^oC$ was slowly added $POCl_3$ (10.0 mL, 107.3

mmol). The reaction was allowed to warm to room temperature. The reaction was quenched with cold water followed by the addition of $NaHCO_3$ (saturated). The resulting crude mixture was extracted with EtOAc and washed with brine, dried over sodium sulfate and concentrated *in vacuo.* The resulting crude product was then separated by column chromatography (Rf =1:4 EtOAc/Hexane) to provide an off white solid (6.16 grams, 24.9 mmol, 28% yield). $^1$H NMR (500 MHz, Chloroform-d) δ 9.67 (s, 1H), 7.55 (dd, J = 8.6, 2.0 Hz, 1H), 7.46 (dd, J = 2.1, 1.0 Hz, 1H), 6.60 (d, J = 8.6 Hz, 1H), 3.71-3.698 (m, 5H), 3.40 (dd, J = 6.6, 4.8 Hz, 3H), 2.77 (t, J = 6.3 Hz, 3H), 2.64 (t, J = 7.2 Hz, 3H), 2.06 – 1.82 (m, 3H), -0.00 (s, 3H). ESI-MS m/z 270.1 $[M+Na]^+$

**Methyl 3-(6-(hydroxymethyl)-3,4-dihydroquinolin-1(2*H*)-yl)propanoate:**

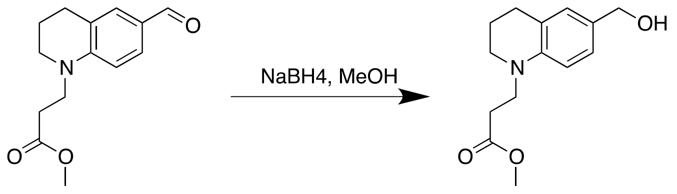

To a stirring solution of substrate (4.34 grams, 17.6 mmol) in MeOH (100 mL) at room temperature was added a tablet of $NaBH_4$ (1 gram, 26.4 mmol). After 35 minutes, an additional tablet of $NaBH_4$ (1 gram, 26.4 mmol) was added. After an additional 45 minutes the reaction was quenched by the addition of acetone. The crude reaction mixture was then concentrated *in vacuo.* The resulting crude mixture was diluted in brine and EtOAc. The product was extracted with EtOAc and washed with brine. The organic layer was dried over sodium sulfate and concentrated *in vacuo.* The resulting pink oil, which degrades quickly, was carried onto the next reaction without further purification (3.96 grams, 15.9 mmol, 91% yield).

**Atto610 Methyl Propanoate:**

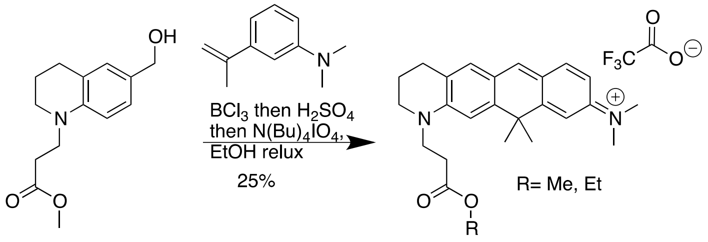

The following procedure is adapted from previously published literature[1,2]. To a stirring solution of substrate (1.97, grams, 7.48 mmol) and *N*,*N*-dimethyl-3-(prop-1-en-2-yl)aniline (1.29 grams, 8.00 mmol) in DCM (16 mL) at 0 °C was added boron trichloride. The reaction was allowed to continue at 0°C for 1 hour. The reaction was then allowed to warm to RT. After an additional hour of reacting, $H_2SO_4$ (40 mL, concentrated) was carefully added and DCM was removed *in vacuo.* This reaction was allowed to continue for an additional 2 hours. A copious amount of ethanol (~ 500 mL) was added followed by the addition of $(nBu)_4NIO_4$ (0.81 grams, 1.80 mmol). The reaction was then refluxed for 10 minutes. The reaction was then allowed to cool to room temperature and stir overnight. The crude mixture was then concentrated *in vacuo* and the product was extracted with several portions of DCM. The crude product was purified by reverse phase HPLC (10-90% ACN/$H_2O$ over 20 minutes) to provide a blue powdery solid TFA salt (close to 1:1 mixture of ethyl ester and methyl ester, 0.926 grams, 1.84 mmol, 25%). For the ethyl ester compound: $^1$H NMR (600 MHz, Acetonitrile-d3) δ 7.89 (s, 1H), 7.69 – 7.53 (m, 1H), 7.30 (s, 1H), 7.11 – 6.98 (m, 1H), 6.84 (dd, J = 9.1, 2.3 Hz, 1H),

4.15 – 3.99 (m, 2H), 3.88 (t, J = 7.0 Hz, 2H), 3.56 (t, J = 5.7 Hz, 2H), 3.23 (d, J = 1.3 Hz, 6H), 2.71 (t, J = 6.3 Hz, 2H), 2.66 (dd, J = 7.6, 6.3 Hz, 2H), 1.61 (d, J = 1.3 Hz, 6H), 1.15 (m, 3H); MS-ESI found m/z 405.3 $[M+H]^+$

**Atto610 Propanoic Acid:**

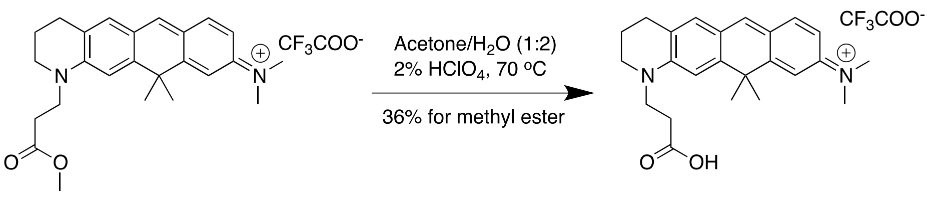

The substrate (600.0 mg, 1.19 mmol) was dissolved in a perchloric acid solution of acetone/water (3 mL: 50 mL: 100 mL, acid/acetone/water) and heated overnight at 70°C under a reflux condenser. The acetone was removed *in vacuo* and the product was extracted with DCM, dried over sodium sulfate and concentrated *in vacuo*. The crude product was purified on the reverse phase HPLC (10-90% ACN/$H_2O$ over 15 minutes). The acetonitrile was removed *in vacuo* and the product was extracted in DCM, dried over sodium sulfate and concentrated *in vacuo* to provide the pure product as a purple, solid TFA salt ( 212.7 mg, 0.434 mmol, 36%) $^1H$ NMR (500 MHz, Acetonitrile-d3) δ 7.93 (s, 1H), 7.63 (d, J = 9.1 Hz, 1H), 7.35 (s, 1H), 7.11 (s, 1H), 7.07 (d, J = 2.2 Hz, 1H), 6.89 (dd, J = 9.1, 2.4 Hz, 1H), 3.93 (t, J = 7.0 Hz, 2H), 3.63 (t, J = 5.7 Hz, 2H), 3.28 (s, 6H), 2.81 – 2.71 (m, 4H), 1.66 (s, 6H). MS-ESI m/z 377.3 $[M+H]^+$

**Atto610ADA—DA:**

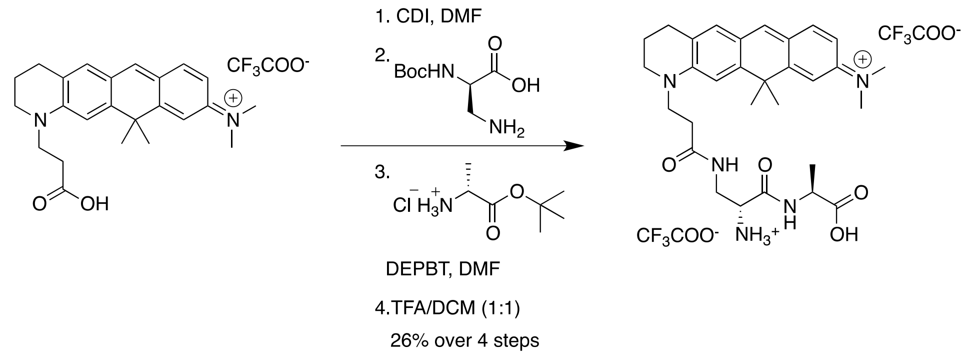

Under a blanket of argon the substrate (41.2 mg, 0.084 mmol) was dissolved in DMF (0.8 mL). To this stirring solution was added CDI (19.9 mg,  0.123) in one portion. After 2.5 hours, Boc-D-2,3-diaminopropionic acid was added (22.5 mg, 0.111 mmol). After 12 hours 1.5 mL of the following mixture was added to the reaction: DMF (1 mL), DEPBT (0.516 mg, 1.7mmol) DIEA (0.58 mL, 3.32 mmol), D-AlaOtBu (482.6 mg, 3.32 mmol). After about one hour of reaction, the reaction mixture was partitioned between DCM and 1 M HCl. The organic phase was saved and dried over Na2SO4. The crude product was concentrated *in vacuo* and diluted in a TFA/DCM mixture (1:1, 20 mL). After deprotection (~ 5 hours) the acidic solvent was removed *in vacuo* and the crude product was isolated off of reverse phase HPLC (10-90% ACN/$H_2O$ over 15 minutes, RT = 9.2 minutes) to provide a beautiful purple, powdery solid (16.6 mg, 26%) $^1$H NMR (500 MHz, Acetonitrile-d3) δ 8.14 (s, 1H), 7.96 (s, 1H), 7.71 (s, 1H), 7.46 (d, J = 9.0 Hz, 1H), 7.10 (s, 1H), 6.94 (s, 1H), 6.87 (d, J = 2.4 Hz, 1H), 6.72 (dd, J = 8.9, 2.3 Hz, 1H), 4.38 – 4.18 (m, 1H), 4.10 (s, 1H), 3.77 (s, 2H), 3.59 (s, 2H), 3.44 (d, J = 5.7 Hz, 2H), 3.12 (s, 6H), 2.55 (t, J = 6.2 Hz, 2H), 2.49 (s, 2H), 1.81 (p, J = 2.5 Hz, 3H), 1.76 (d, J = 8.0 Hz, 2H Note: NH3, disappears when D2O is added), 1.45 (s, 6H), 1.24 (d, J = 7.1 Hz, 3H); $^{13}$C NMR (126 MHz, cd3cn) δ 174.71, 173.92, 167.75, 157.84, 157.47, 157.44, 155.22, 153.79, 139.17, 137.15, 125.32, 121.90, 121.37, 118.26, 113.62, 111.87, 111.53, 54.49,

51.68, 49.84, 49.08, 47.17, 42.60, 41.26, 41.13, 33.89, 33.76, 27.39, 21.50, 17.33, 8.96; HRMS-ESI-TOF m/z calc 534.0380, Found 534.0386 $[M+H]^+$

**FDAAD labeling conditions.** Growth and labeling conditions for strains used in supplementary figures 2C and 2D are shown below.

| **Figure** | **Bacteria** | **Growth condition** | **Label** |
|---|---|---|---|
| Supplementary Figure 2 C | *E. coli* | M9 + 0.2% glucose, 37ºC | 3 mM NADA—DA, Overnight |
| | *E. coli* | M9 + 0.2% glucose, 37ºC | 3 mM DA—NADA, Overnight |
| | *B. subtilis ΔdacA* | SSM + 1% LB, 37ºC | 5 mM NADA—DA, 90 min |
| | *B. subtilis ΔdacA* | SSM + 1% LB, 37ºC | 5 mM DA—NADA, 90 min |
| | *S. venezuelae* | LB, 30ºC | 2 mM NADA—DA, 12 min |
| | *S.* venezuelae | LB, 30ºC | 2 mM DA—NADA, 12 min |
| Supplementary Figure 2 D | *E. coli* | LB, 37ºC | 2 mM Atto$_{610}$ADA—DA, 12 min |
| | *E. coli* | M9 + 0.2% glucose, 37ºC | 2 mM BADA—DA, 12 min |
| | *E. coli* | LB, 37ºC | 2 mM BADA—DA, 12 min |
| | *B. subtilis* | S750 + 1% glucose, 37ºC | 1 mM Atto$_{610}$ADA—DA, 30 min |
| | *B. subtilis* | S750 + 1% glucose, 37ºC | 0.5 mM BADA—DA, 40 min |

### SUPPLEMENTARY TABLES

**Supplementary Table 1.** Bacterial strains and plasmids used in this study.

| Strain or Plasmid | Relevant Genotype, Features or Characteristics | Source or Reference |
|---|---|---|
| **Source Plasmids** | | |
| pNTPS139 | Kmr; Suicide vector containing oriT and sacB | D. Alley |
| pUC18-mini-Tn7T-GM-Plac | $Ap^r$ $Gm^r$; mini-Tn7 vector containing *lacI*$^q$ and lac promoter | Figueroa-Cuilan et al[1] |
| pTNS3 | $Ap^r$; helper plasmid encoding the site-specific TnsABCD Tn*7* transposition pathway | Choi et al[2] |
| pGCH14 | pG18mob carrying the LacI-repressible *repABC* operon, $Gm^r$ | Krol et al[3] |
| pK18mobsacB | suicide vector, *lacZ*, *mob*, *sacB,* $Km^r$ | Schäfer et al[4] |
| pSRKKm | pBBR1MCS-5-derived broad-host-range expression vector containing *lac* promoter and *lacI*$^q$, *lacZα*$^+$, $Km^r$ | Khan et al[5] |
| **Deletion Plasmids** | | |
| pNTPS139Δ*pbp1a* | $Km^r$ $Suc^s$; deletion plasmid for *pbp1a* | This Study |
| pNTPS139Δ*pbp1b1* | $Km^r$ $Suc^s$; deletion plasmid for *pbp1b1* | This Study |
| pNTPS139Δ*pbp1b2* | $Km^r$ $Suc^s$; deletion plasmid for *pbp1b2* | This Study |
| pNTPS139Δ*pbp1c* | $Km^r$ $Suc^s$; deletion plasmid for *pbp1c* | This Study |
| pNTPS139Δ*mtgA* | $Km^r$ $Suc^s$; deletion plasmid for *mtgA* | This Study |
| pNTPS139Δ*pbp3a* | $Km^r$ $Suc^s$; deletion plasmid for *pbp3a* | This Study |
| pNTPS139Δ*pbp3b* | $Km^r$ $Suc^s$; deletion plasmid for *pbp3b* | This Study |
| pK18mobsacΔ*mrcA1* | $Km^r$ $Suc^s$; deletion plasmid for *mrcA1* | This Study |

| | | |
|---|---|---|
| pK18mobsacΔ*mrcA2* | $Km^r$ $Suc^s$; deletion plasmid for *mrcA2* | This Study |
| pK18mobsacΔ*mrcB* | $Km^r$ $Suc^s$; deletion plasmid for *mrcB* | This Study |
| pK18mobsacΔ*pbp* | $Km^r$ $Suc^s$; deletion plasmid for *pbp* | This Study |
| pK18mobsacΔ*pbpC* | $Km^r$ $Suc^s$; deletion plasmid for *pbpC* | This Study |
| pK18mobsacΔs*mc02856* | $Km^r$ $Suc^s$; deletion plasmid for *smc02856* | This Study |
| **Complementation Plasmids** | | |
| pGCH14-mrcA1 | $Gm^r$ | This Study |
| **Depletion Plasmids** | | |
| pUC18-mini-Tn7T-GM-plac pbp1a | $Ap^r$ $Gm^r$; mini-Tn7 vector containing *pbp1a* under control of the lac promoter | This Study |
| pUC18-mini-Tn7T-GM-plac pbp3a | $Ap^r$ $Gm^r$; mini-Tn7 vector containing *pbp3a* under control of the lac promoter | This Study |
| ***E. coli* strains** | | |
| DH5α | Cloning strain | Life Technologies |
| S17-1 | Smr;RP4-2 TC::MU Km-Tn7; for plasmid mobilization | Simon et al[6] |
| MG1655 | Strain used for labeling studies | P. Foster – Indiana University |
| ***A. tumefaciens* strains** | | |
| C58 | Parent strain | Watson et al[7] |
| C58ΔtetRA::a-attTn7 (WT) | Replacement of the tetRA locus with an artificial attTn7 site | Figueroa-Cuilan et al[1] |
| C58ΔtetRA::a-attTn7 Δ*pbp3a* | Δ*pbp3a* | This Study |
| C58ΔtetRA::a-attTn7 Δ*pbp3b* | Δ*pbp3b* | This Study |
| C58ΔtetRA::a-attTn7 Δ*pbp1b1* | Δ*pbp1b1* | This Study |
| C58ΔtetRA::a-attTn7 Δ*pbp1b2* | Δ*pbp1b2* | This Study |
| C58ΔtetRA::a-attTn7 Δ*pbp1c* | Δ*pbp1c* | This Study |
| C58ΔtetRA::a-attTn7 Δ*mtgA* | Δ*mtgA* | This Study |
| C58ΔtetRA::a-attTn7 Δ*pbp1b1*,Δ*pbp1b2* | Δ*pbp1b1*,Δ*pbp1b2* | This Study |

| | | |
|---|---|---|
| C58ΔtetRA::a-attTn7 Δ*pbp1b1, Δpbp1b2, Δpbp1c* | Δ*pbp1b1,*Δ*pbp1b2,*Δ*pbp1c* (Δ3pbp*)* | This Study |
| C58ΔtetRA::mini-Tn7-GM-Plac-pbp1a | Mini-Tn7T-GM-Plac-pbp1a inserted into a-attTn7 site | This Study |
| C58ΔtetRA::mini-Tn7-GM-Plac -pbp1a, Δ*pbp1a* | Chromosome-based complementation of Δ*pbp1a* with C58ΔtetRA::mini-Tn7-GM-Plac-pbp1a allowing depletion of PBP1a under control of the lac promoter | This Study |
| C58ΔtetRA::mini-Tn7-GM-Plac-pbp3a | Mini-Tn7T-GM-Plac-pbp3a inserted into a-attTn7 site | This Study |
| C58ΔtetRA::mini-Tn7-GM-Plac-pbp3a, Δ*pbp3a* | Chromosome-based complementation of Δ*pbp3a* with C58ΔtetRA::mini-Tn7-GM-Plac-pbp3a allowing depletion of PBP3a under control of the lac promoter | This Study |
| C58ΔtetRA::mini-Tn7-GM-Plac-pbp3a, Δ*pbp3a,* Δ*pbp3b* | Chromosome-based complementation of Δ*pbp3a* with C58ΔtetRA::mini-Tn7-GM-Plac-pbp3a allowing depletion of PBP3a under control of the lac promoter, Δ*pbp3b* | This Study |
| ***S. meliloti* strains** | | |
| Rm2011 | Wild type, $Str^r$ | Casse et al[8] |
| Rm2011 *rgsP-egfp* | Rm2011 expressing *egfp*-tagged *rgsP*, markerless insertion | Schäper et al[9] |
| Rm2011 rgsP-*egfp*, Δ5pbp | Rm2011 rgsP-egfp carrying markerless deletions of mrcA2, mcrB, pbp, pbpC and SMc02856 Δ5pbp | This Study |
| Rm2011 rgsP-egfp mrcA1 depletion | Rm2011 rgsP-egfp carrying markerless deletion of mrcA1, curable complementation plasmid pGCH14-mrcA1, and pSRKKm as a source of lacI to cure pGCH14-mrcA1, $Gm^r$ $Km^r$ | This Study |
| ***B. abortus* strain** | | |
| S19 | Wild type | J. Skyberg – University of Missouri |
| ***B. subtilis* strains** | | |

| | | |
|---|---|---|
| PY76 | Wild type | D. Kearns – Indiana University |
| PY76, *dacA::cam* | *ΔdacA* | D. Kearns – Indiana University |
| ***S. venezuelae* strain** | | |
| *S. venezuelae* | Wild type | J. Nodwell – McMaster University |

[1]Figueroa-Cuilan W, Daniel JJ, Howell M, Sulaiman A, Brown PJ. Mini-Tn7 Insertion in an Artificial *att*Tn7 Site Enables Depletion of the Essential Master Regulator CtrA in the Phytopathogen *Agrobacterium tumefaciens*. Appl Environ Microbiol. 2016. 82:5015-25.

[2]Choi KH, Mima T, Casart Y, Rholl D, Kumar A, Beacham IR, Schweizer HP. Genetic tools for select-agent-compliant manipulation of *Burkholderia pseudomallei*. Appl Environ Microbiol. 2008. 74:1064-75.

[3]Krol E, Yau HCL, Lechner M, Schäper S, Bange G, Vollmer W, Becker A. Tol-Pal System and Rgs Proteins Interact to Promote Unipolar Growth and Cell Division in *Sinorhizobium meliloti*. mBio. 2020. 11:e00306-20.

[4]Schäfer A, Tauch A, Jäger W, Kalinowski J, Thierbach G, Pühler A. 1994. Small mobilizable multi-purpose cloning vectors derived from the *Escherichia coli* plasmids pK18 and pK19: selection of defined deletions in the chromosome of *Corynebacterium glutamicum*. Gene. 1994. 145:69-73.

[5]Khan SR, Gaines J, Roop RM, Farrand SK. Broad-host-range expression vectors with tightly regulated promoters and their use to examine the influence of TraR and TraM expression on Ti plasmid quorum sensing. Appl Environ Microbiol. 2008. 74:5053-5062.

[6]Simon R, Priefer U, Pühler A. A broad host range mobilization system for *in vivo* genetic engineering: transposon mutagenesis in Gram-negative bacteria. Nature Biotechnol. 1983. 1:784-791.

[7]Watson B, Currier TC, Gordon MP, Chilton MD, Nester EW. Plasmid required for virulence of *Agrobacterium tumefaciens*. J Bacteriol. 1975. 123:255-264.

[8]Casse F, Boucher C, Julliot J, Michel M, Dénarié J. Identification and characterization of large plasmids in *Rhizobium meliloti* using agarose gel electrophoresis. Microbiology. 1979. 113:229-242.

[9]Schäper S, Yau HCL, Krol E, Skotnicka D, Heimerl T, Gray J, Kaever V, Søgaard-Andersen L, Vollmer W, Becker A. Seven-transmembrane receptor protein RgsP and cell wall-binding protein RgsM promote unipolar growth in Rhizobiales. PLoS Genet. 2018. 14:e1007594.

### **Supplementary Table 2.** Synthesized DNA primers used in this study

| **Synthesized DNA** | **Sequence (5' – 3')** |
|---|---|
| **Primers for gene amplification in *A. tumefaciens*** | |
| PBP1a For NdeI | `CGCGATCATATGATCAGACTGATTGGA` |
| PBP1a Rev BamHI stop codon | `GCTCACGGATCCTCAATAAAGACCGCCGCCAC` |

| | |
|---|---|
| PBP3a For NdeI | CGCGATCATATGTCTTTCCTTTCCCGT |
| PBP3a Rev BamHI stop codon | CGCTGGATCCTCAATAAGACACGAGCAAG |
| **Primers for deletion vectors in *A. tumefaciens*** | |
| PBP1a P1 For SpeI | GCACACTAGTTTATGCCGGTTTCATGGTTCTCCG |
| PBP1a P2 Rev | AAGCTTGGTACCGAATTCACCAAGCTACCGATAATTCGA |
| PBP1a P3 For | GAATTCGGTACCAAGCTTCATCAGTCATGACGTTTGGCG |
| PBP1a P4 Rev BamHI | CTAGGGATCCGCGCCGGAATGCACTTCCACATAG |
| PBP1a P5 For | CTGAAGCAGAAGGGAATTC |
| PBP1a P6 Rev | GGAAGAAAACGAGGTGTGAC |
| PBP1b1 P1 For SpeI | GCATACTAGTTCGGCGACGGTTGCACTGGCCGCT |
| PBP1b1 P2 Rev | AAGCTTGGTACCGAATTCCCAGCCTAAGTCGCTCCTATT |
| PBP1b1 P3 For | GAATTCGGTACCAAGCTTCCCGCGTCTGGTAATGGGCCA |
| PBP1b1 P4 Rev BamHI | GTACGGATCCAATGCGACGGTCGCCAATACAGGG |
| PBP1b1 P5 For | GTATTGTCAGTCCAATCGG |
| PBP1b1 P6 Rev | TGGGCCCACCAGCGACATG |
| PBP1b2 P1 For SpeI | GCACACTAGTCACAAGCATGCCTAGGTTTTGCGTCGG |
| PBP1b2 P2 Rev | AAGCTTGGTACCGAATTCGAAATGATCAGGCATCTGGTC |
| PBP1b2 P3 For | GAATTCGGTACCAAGCTTGTGACCATTCCGGTGATG |
| PBP1b2 P4 RevBamHI | CTAGGGATCCCTAACGCCGCCCCGCTTC |
| PBP1b2 P5 For | GCGGTTCTCGTAGTCGGAG |
| PBP1b2 P6 Rev | GTGGAAAGAATATTCGGC |
| PBP1c P1 For SpeI | GTATACTAGTCCGGCGCAGCCGCTTGCCGCCGGT |
| PBP1c P2 Rev | AAGCTTGGTACCGAATTCGATGCCGGCGATGACAGCCTT |
| PBP1c P3 For | GAATTCGGTACCAAGCTTGGTCTGCCACCGAAACGCCAA |
| PBP1c P4 Rev BamHI | GCAGGGATCCATTGCCGTGACGGAACATT |
| PBP1c P5 For | GCTTGCCGGTGCCGTGGCG |
| PBP1c P6 Rev | TATCATGTCCGACACCGATG |
| PBP3a P1 For SpeI | GCAGACTAGTATCAAGCACAAGGCCGATCTGAAG |

| | |
|---|---|
| PBP3a P2 Rev | AAGCTTGGTACCGAATTCCAGCGGTTCACCGATCCTGTT |
| PBP3a P3 For | GAATTCGGTACCAAGCTTTTGTTAGCTTATGATGTTCCG |
| PBP3a P4 Rev BamHI | GCTGGGATCCTAATCTTCGCGACCAGGAGCGATG |
| PBP3a P5 For | GGACAAGCCCGCCATTTTC |
| PBP3a P6 Rev | AGCGCCTTTTTCCATCAGC |
| PBP3b P1 For SpeI | GCAGACTAGTGACCTGGGCGCAGCTCGGCCGCCA |
| PBP3b P2 Rev | AAGCTTGGTACCGAATTCGTCTGATGCCGCCCACTTCAT |
| PBP3b P3 For | GAATTCGGTACCAAGCTTGCCCTGACTTGGTGAGGGAGG |
| PBP3b P4 BamHI | CTCGGGATCCTTGCGATCCTCACCTGGCATGCGG |
| PBP3b P5 For | ACGGCGGCGGCACCAAGGG |
| PBP3b P6 Rev | CTCGGGCGCACGGCGGAAA |
| MtgA P1 For SpeI | GCACACTAGTAGGCGATTATGTCGAAAGC |
| MtgA P2 Rev | AAGCTTGGTACCGAATTCTGCCGTCTTCAAGGC |
| MtgA P3 For | GAATTCGGTACCAAGCTTTCCTGCGTGCTTGACTG |
| MtgA P4 Rev BamHI | CTAGGGATCCGCACTTGGCCATGAGATC |
| MtgA P5 For | CGAAGAGGCGCAGTC |
| MtgA P6 Rev | CATCTGCACGGCGGCAGC |
| **Primers for deletion and depletion vectors in *S. meliloti*** | |
| MrcA1 500up For XbaI | ctgtctagaGCTCGAGCTCGGTTTCCTGTC |
| MrcA1 stop codon Rev NcoI | atatccatggTCAGAACAGTCCGTTGGAGCC |
| MrcA1 P1 For HindIII | atataagcttATCTTGCCGCACGCGAGAAT |
| MrcA1 P2 Rev XbaI | atattctagaCAGTCAGGTACCGGTATCTA |
| MrcA1 P3 For XbaI | atattctagaACCTCCGGCTCCAACGGACT |
| MrcA1 P4 Rev EcoRI | atatgaattcGACGTATGACGGCGAGCGTT |
| MrcA2 P1 For HindIII | atataagcttACAGGTGCATTGCGTCTGTG |
| MrcA2 P2 Rev XbaI | atattctagaGCTGATACCAATGGTTCAGAC |
| MrcA2 P3 For XbaI | atattctagaTGGCGATGAGCGGCAGACCT |
| MrcA2 P4 Rev EcoRI | atatgaattcCGATAGTCATGGATGCGTTG |

| | |
|---|---|
| MrcB P1 For PstI | atatctgcagCCACATTGCGGACAGTACAG |
| MrcB P2 Rev XbaI | atattctagaTGGTTACCGTAAAAGGCTCC |
| MrcB P3 For XbaI | atattctagaTGCCTGGACTACCGACTCAG |
| MrcB P4 Rev EcoRI | atatgaattcCGGTCGGCACTAGCCGCGAA |
| PBP P1 For HindIII | atataagcttGACGACGAACTCGTTCGCCA |
| PBP P2 Rev XbaI | atattctagaATGACGATATTTGCACGGGG |
| PBP P3 For XbaI | atattctagaCTTTGCAGTCAGAAATCCGT |
| PBP P4 Rev EcoRI | atatgaattcCAGGGCGAGTATGATCACGA |
| PBPC P1 For HindIII | atataagcttATCATTCCCGAGCTATCCAG |
| PBPC P2 Rev XbaI | atattctagaGCGTCCTACTGGGCTGCCTG |
| PBPC P3 For XbaI | atattctagaCCAGCGTTCGCGTCTTCGTC |
| PBPC P4 Rev EcoRI | atatgaattcTCCTTGAGCAACATGGACGA |
| Smc02856 P1 For EcoRI | tatagaattcGACGATGGTGAAGGACGTG |
| Smc02856 P2 Rev XbaI | atatctagaCATCGCCCGCAAGCGATCC |
| Smc02856 P3 For XbaI | atatctagaCTGTTCGACCTGCTGACCGG |
| Smc02856 P4 Rev HindIII | atataagcttATGCGCAGCACGTCGAGACC |
